# A Bioactive Phospholipid Promotes Rapid Lung Progenitor Activation via AP-1

**DOI:** 10.64898/2026.07.28.741214

**Authors:** Alena Klochkova, Aaron I. Weiner, Kentaro Hara, Nicolas P. Holcomb, Sara Kass-Gergi, Meryl Mendoza, Evelyn A. Martinez, Diana M. Abraham, Melanie Y. Staszewsli, Michael M. Maiden, Jeremy Katzen, Andrew E. Vaughan

## Abstract

Upon injury to the distal lung, alveolar type 2 cells (AT2s) must make a discrete switch from surfactant factories to stem cells capable of regeneration, which involves both proliferation and differentiation into oxygen-exchanging alveolar type 1 (AT1) cells. However, the discrete signals and molecular pathways facilitating this fundamental switch in AT2 functionality are uncertain. Here we demonstrate that the bioactive lipid lysophosphatidic acid (LPA), typically associated with driving fibrosis, is an extremely efficient inducer of this state change in comparison to previously implicated signals IL-1β and p53 stabilization. We observed endogenous production and accumulation of LPA in influenza-injured murine lungs, creating a microenvironment that facilitates AT2 progenitor switching. Multiple transcriptomic approaches reveal elevation of Fosl1 and Jun, core members of the Activator Protein 1 (AP-1) transcription factor family, in response to LPA. Using novel genetic models combined with influenza injury, we demonstrate that AP-1 activity in AT2s is necessary for effective alveolar regeneration at both the cellular and physiologic levels. These findings unveil a critical relationship between paracrine LPA and cell-intrinsic AP-1 in facilitating effective lung alveolar regeneration.

**Highlights:**

1. LPA facilitates progenitor switching in lung regeneration via initiation of a discrete transcriptomic state
2. LPA promotes AT2 state switching via Jun (AP-1)
3. Impaired AP-1 signaling significantly restricts recovery from influenza infection

## Introduction

Mammalian lungs require a massive alveolar surface area to facilitate efficient gas exchange. However, this results in ample exposure of the lung epithelium to numerous environmental hazards and pathogenic microorganisms. This intrinsic vulnerability to damage necessitates a robust capacity for repair to maintain pulmonary function. Alveolar type 2 (AT2) cells are key players in this epithelial regenerative capacity, where they act as facultative (non-professional) stem cells.^1^ While serving a critical homeostatic role of surfactant synthesis, upon injury AT2s can proliferate and differentiate into oxygen-exchanging AT1 cells^1,2^. AT2 cells’ ability to restore the physiologic cellular structure of the alveoli is a crucial component of effective, euplastic lung regeneration. While the distinct homeostatic vs regenerative functions of AT2s are both very well described,^3,4^ *how* AT2 cells “toggle” between these two very different functional roles remains largely unknown.

Recent lineage-tracing and transcriptomics studies revealed that upon injury, AT2 cells can undergo a transition into a unique cellular state characterized by high expression of *Krt8* ^5–8^ and additional more specific markers including *Cldn4* and *Lgals2*.^4,8^ Despite shared recognition that these “transitional” cells (the term we will primarily use in this study) emerge during injury, the physiologic or pathophysiologic function of transitional cells remain hotly debated. Some studies suggest this emergent state is explicitly pathological and, via secretion of paracrine factors, pro-fibrotic.^5^ Other studies however point to the transitional state as representing an intermediate stage through which AT2 cells must traverse in order to differentiate into AT1s.^8^ Our studies here suggest an additional scenario: that the transitional state represents cells actively undergoing the “toggle / switch” between surfactant factories and dedicated stem cells.

In the present study we explore how microenvironmental and cell-intrinsic signals drive initiation of this distinct AT2 stem cell state to promote epithelial regeneration in response to influenza injury. Critically, and surprisingly given its typical identification as a pro-fibrotic factor, we demonstrate a key role for the bioactive phospholipid signaling molecule lysophosphatidic acid (LPA) in promoting the AT2 switch into the stem cell state. We observe that LPA promotes a much more rapid and robust switch than previously implicated signals, and that this occurs in part by engagement of Activator Protein-1 (AP-1) family genes. We subsequently employ a newly generated transgenic animal model to demonstrate that AP-1 is absolutely necessary for the successful engagement of AT2 stem cell functionality *in vivo* and *in vitro* at the cellular level, and that by blocking AP-1, mice are unable to functionally regenerate after influenza injury. These findings uncover critical external and cell-intrinsic players necessary for AT2 cells to fully engage in their remarkable functionality as stem cells of the delicate alveolar compartment.

## Results

### Lysophosphatidic acid is a potent mediator of AT2 phenotypic switching

Though the pro-fibrotic effects of lysophosphatidic acid on lung fibroblasts are well known in the context of pulmonary fibrosis,^9–11^ LPA also has notable protective and regenerative effects in other contexts.^12,13^ As such, we tested whether LPA has similar effects on the predominant stem cell population of the alveoli, AT2s. Given that IL-1β signaling and p53 stabilization have both been implicated in adoption of AT2 progenitor properties,^6–8^ we directly compared the effects of LPA stimulation to these established factors (**Figure 1A**). We first isolated primary AT2 cells by fluorescence-activated cell sorting (**Figure S1**) using a well-validated gating strategy.^14^ Isolated AT2s were seeded on matrigel and cultured as organoids for 9 days. Organoids were then “starved” for 22 hours followed by 6 hours treatment with IL-1β, Nutlin-3a, 1-Olyeyl-LPA, or a vehicle control (**Figure 1B**). We then proceeded with bulk RNA-Seq (**Figure S1**). The effect of each factor was first validated by examining the known target genes (**Figure 1C, 1D, 1E** (*top*)). In agreement with published literature,^15^ IL-1β successfully induced expression of *Cxcl1* and *Cxcl2* (**Figure 1C** (*top*)), and *Trmp4* was downregulated by Nutlin-3a (**Figure 1D** (*top*)) as expected.^16^ Upregulation of *Ptgs2* was confirmed as a control for LPA treatment (**Figure 1E** (*top*)).^17^

**Figure 1.**
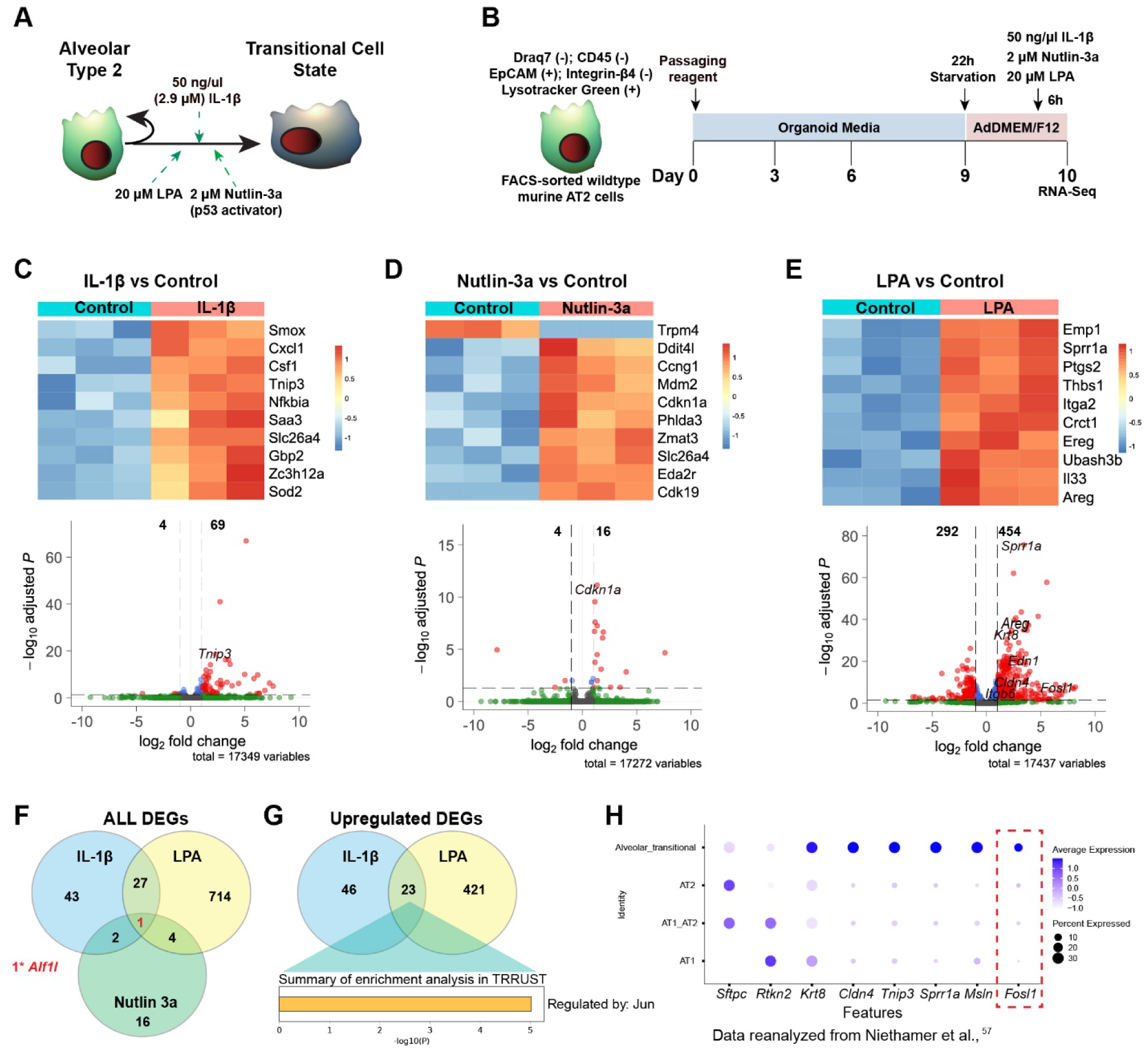
Lysophosphatidic acid drives rapid activation of the transitional cell state. **(A)** Model of AT2 cells proliferation and differentiation via transitional cell state highlighting tested signaling molecules. **(B)** Experimental outline for stimulation of primary mouse AT2s. CD45-, draq7-, EpCAM+, Integrin-β4-, Lysotracker green+ single cells were isolated by FACS from 3 C57BL/6 mice (10 weeks-old 2 female, 1 male) and plated on the top of matrigel (7000-15000 cells per well). On day 9 the cells were starved for 22 hours and then treated for 6 hours with 50 ng/µl IL-1β, 2 µM Nutlin-3a, 20 µM LPA or vehicle control as presented in the schema design. On day 10, RNA was isolated from organoids and used for bulk RNA-seq. **(C)** Heatmap for top 10 differentially expressed genes (DEGs) as determined by DESeq2 (*top*) after 6h IL-1β treatment as well as volcano plot (*bottom*) with selected transitional state-associated genes. **(D)** Heatmap for top 10 DEGs as determined by DESeq2 (*top*) after 6h Nutlin-3a treatment as well as volcano plot (*bottom*) with selected transitional state-associated genes. **(E)** Heatmap for top 10 DEGs as determined by DESeq2 (*top*) after 6h LPA treatment as well as volcano plot (*bottom*) with selected transitional state-associated genes **(F)** Venn diagram for all DEGs showing relatively little DEG overlap between these signals except for a single gene *Alf1l*. **(G)** Venn diagram for upregulated DEGs and overlap between IL-1β-and LPA-treated groups. Prediction analysis for transcriptional factors driving the overlapping genes upregulated by IL-1β and LPA identified only *Jun*. **(H)** Dot plot of key genes expression associated with transitional cell state by cell type highlights enrichment of AP-1 family gene *Fosl1* uniquely in transitional cells. Reanalysis of publicly available data.^57^

We observed dramatic changes in LPA-treated cells, identifying >700 differentially expressed genes (DEGs). In sharp contrast, IL-1β induced only 73 DEGs and p53 stabilization with Nutlin-3a induced only 23 DEGs (**Figure 1E, 1C, 1D** (*bottom*)). Strikingly, the majority of the canonical marker genes associated with the transitional cell state were strongly induced by LPA, whereas IL-1β and Nutlin-3a induced only one marker gene each (*Tnip3* and *Cdkn1a*, respectively). Surprisingly, when we analyzed the shared DEGs between all 3 treatments, only one gene, *Alf1l*, was shared (**Figure 1F**). However, analysis of the 23 upregulated DEGs shared between LPA and IL-1β by enrichment analysis in Metascape revealed the <u>only</u> transcription factor predicted to regulate these genes is *Jun*, a core member of AP-1 family (**Figure 1G**). Remarkably, another core AP-1 family member *Fosl1* is highly expressed strictly in transitional state cells but not in AT1s or AT2s (**Figure 1H**), and *Fosl1* was in turn greatly upregulated by LPA, representing the most highly upregulated transcription factor in this dataset (**Figure 1E** (*top*)). Taken together, LPA initiates much more rapid and robust changes in the AT2 transcriptome compared to IL-1β and p53, including very strong induction of nearly all transitional state genes.

### LPA promotes AT2 self-renewal in a distinct progenitor state

To better understand the effects of LPA on AT2s we repeated the transcriptomic analysis, adding more mice per group, testing longer LPA stimulation, and performing deeper sequencing (**Figure 2A**). Notably, these two completely independent experiments recapitulated the core gene expression changes (**Figure 1E**, **Figure 2D,E**), observing >1500 DEGs (larger number here due to higher *n* and deeper sequencing) after a short LPA stimulation, including strong and significant induction of canonical transitional state genes. *Fosl1* was, again, the most highly upregulated transcription factor, which we validated by RT-qPCR (**Figure 2F**).

**Figure 2.**
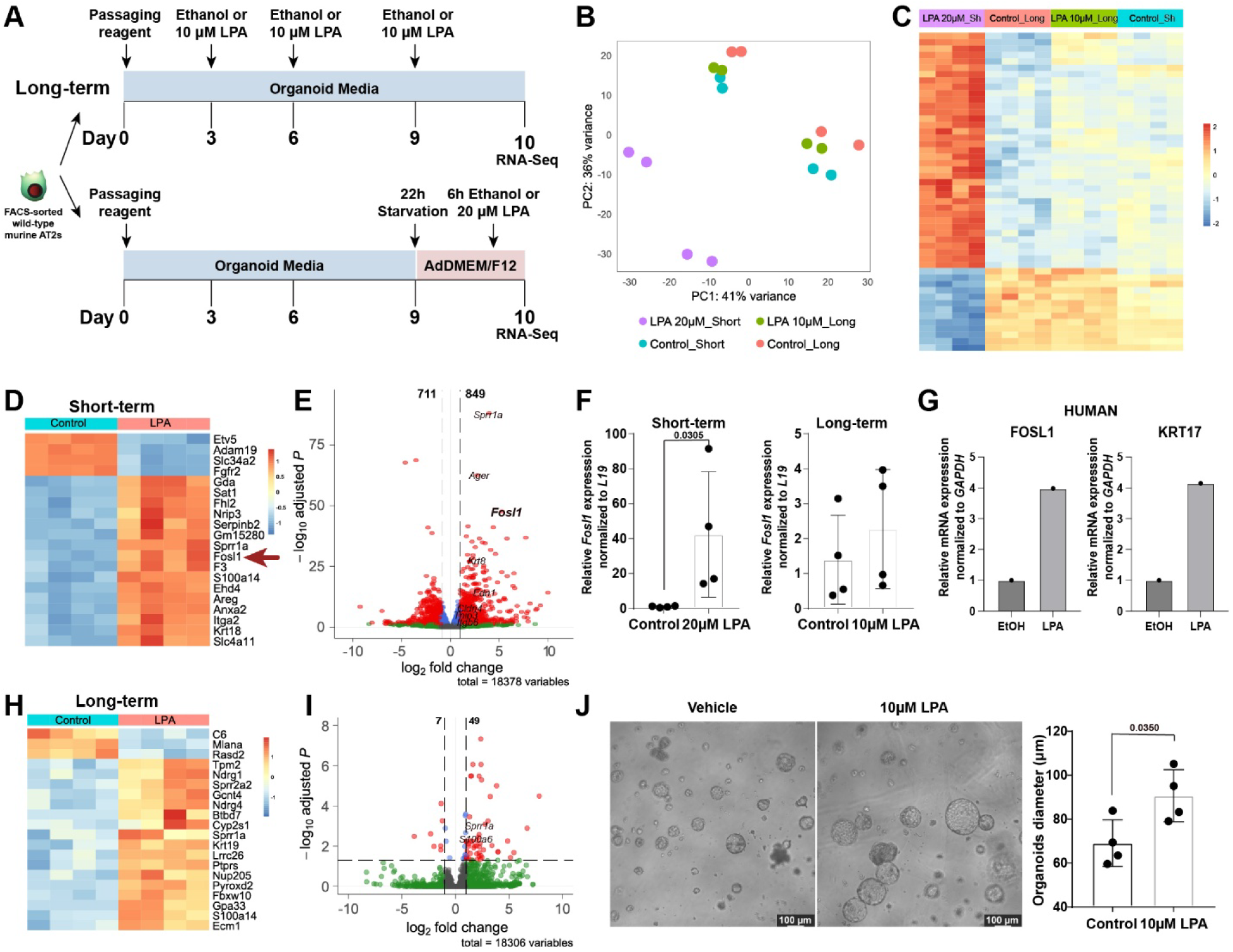
Lysophosphatidic acid promotes broad transcriptional changes indicative of AT2 progenitor switching and self-renewal. **(A)** Experimental design. Freshly derived AT2s were isolated from C57BL/6 mice (8 weeks old, 2 males, 2 female) and plated on the top of matrigel (6000 cells per well). Cells were treated as presented in the schema. For short-term stimulation, cells were treated with 20 µM LPA for 6 hours after 22 hours of starvation or with ethanol as vehicle control. For long-term groups organoids were treated with 10 µM LPA (long-term on the right side) or ethanol as vehicle control through day 10. At day 10, RNA was isolated from organoids for bulk RNA-seq. **(B)** PCA plot demonstrating clustering of samples based on treatment conditions. **(C)** Heatmap highlighting robust transcriptional changes in the short-term LPA treatment group compared to other groups. **(D)** Heatmap for top 20 DEGs as determined by DESeq2 after 6h LPA treatment. **(E)** Volcano plot for short-term treatment with selected transitional state-associated genes. **(F)** *Fosl1* expression in the short-term (*left*) and long-term (*right*) groups determined by RT-qPCR in murine AT2 organoids normalized to its respective vehicle control and *L19*. Data shown as mean fold change ± SD. Each dot represents one biological replicate (n=4). P values were calculated using a one-way ANOVA with post-hoc Turkey multiple comparison test. **(G)** *FOSL1* and *KRT17* expression in human AT2 organoids normalized to *GAPDH,* (n=1). **(H)** Heatmap and **(I)** volcano plot for long-term groups: LPA treatment compared to control. **(J)** Representative light microscopy images of AT2 organoids from long-term experiment, analyzed at day 8 (*left*). Scale bars, 100 µm. Bar graphs represent average organoid diameter (*right*). Each dot represents one biological replicate (n=4). Data shown as mean ± SD (n=4). P value was calculated using a two-tailed unpaired t-test.

LPA was previously reported to be involved in regeneration in other tissues.^12,13^ Specifically, it protects against radiation and chemical injury in the gut^18^ and is also thought to be important in hematopoietic stem cell niche maintenance^19,20^ and hair follicle and tooth development.^21^ We therefore examined the effects of LPA on AT2 self-renewal. We cultured primary murine AT2 cells in presence of 10 µM LPA for 10 days and compared transcriptional changes to short term LPA stimulation above (**Figure 2A**). Interestingly, long-term treatment had a much less dramatic effect (56 DEGs) (**Figure 2B,C**), with only two of the transitional state markers (*Sprr1a* and *S100a14*) retaining significant upregulation (**Figure 2H,I**), though we note most of the transitional genes still trended toward upregulation.

Importantly, despite the more muted transcriptomic changes, we demonstrated that long-term treatment of primary AT2s with LPA *in vitro* significantly increases the size of AT2-derived organoids (**Figure 2J**), promoting self-renewal of AT2s. To address whether the effects of LPA are conserved in human AT2 cells, we repeated the short LPA treatment on established human AT2 organoids as above. In keeping with our murine results, LPA induced expression of *Fosl1* and *Krt17*, the latter a marker of human transitional state cells (**Figure 2G**). Taken together, LPA rapidly and robustly promotes strong changes in the AT2 transcriptome whereas long term LPA stimulation promotes AT2 organoid growth. These findings suggest that LPA treatment initiates an AT2 “progenitor switch”, promoting both adoption of the transitional state and cell proliferation.

### Influenza injury promotes LPA production in damaged lungs

AT2 adoption of the transitional cell state has been previously reported in injured lungs and was attributed to IL-1β and/or p53.^5,6,8^ Because we observed much stronger responses to LPA than either of these signals, we next evaluated whether LPA or LPA-producing enzymes were induced in influenza-injured lungs. Indeed, spatial transcriptomic analysis of murine lungs at 21 days after infection with H1N1 influenza (PR8 strain) (GSE326788)^22^ revealed upregulation of key LPA-producing enzymes including Autotaxin (encoded by *Atx*), Phospholipase A1 (encoded by *Pla1a*), Phospholipase D1 (encoded by *Pld1*) compared to healthy lung (**Figure 3A**). We then analyzed murine lung tissue at 14 days post-infection (dpi) to measure the level of the LPA phospholipid itself (**Figure 3B)**. Visually damaged and adjacent areas of tissue were manually dissected for more granular analysis. We observed significantly increased LPA levels in all damaged tissue at 14 dpi by ELISA (**Figure 3C),** with slightly higher levels in the regions characterized by more obvious tissue damage. Thus, a bioactive lipid with strong transformative effects on AT2s is greatly induced in injured tissue where tissue regeneration is needed.

**Figure 3.**
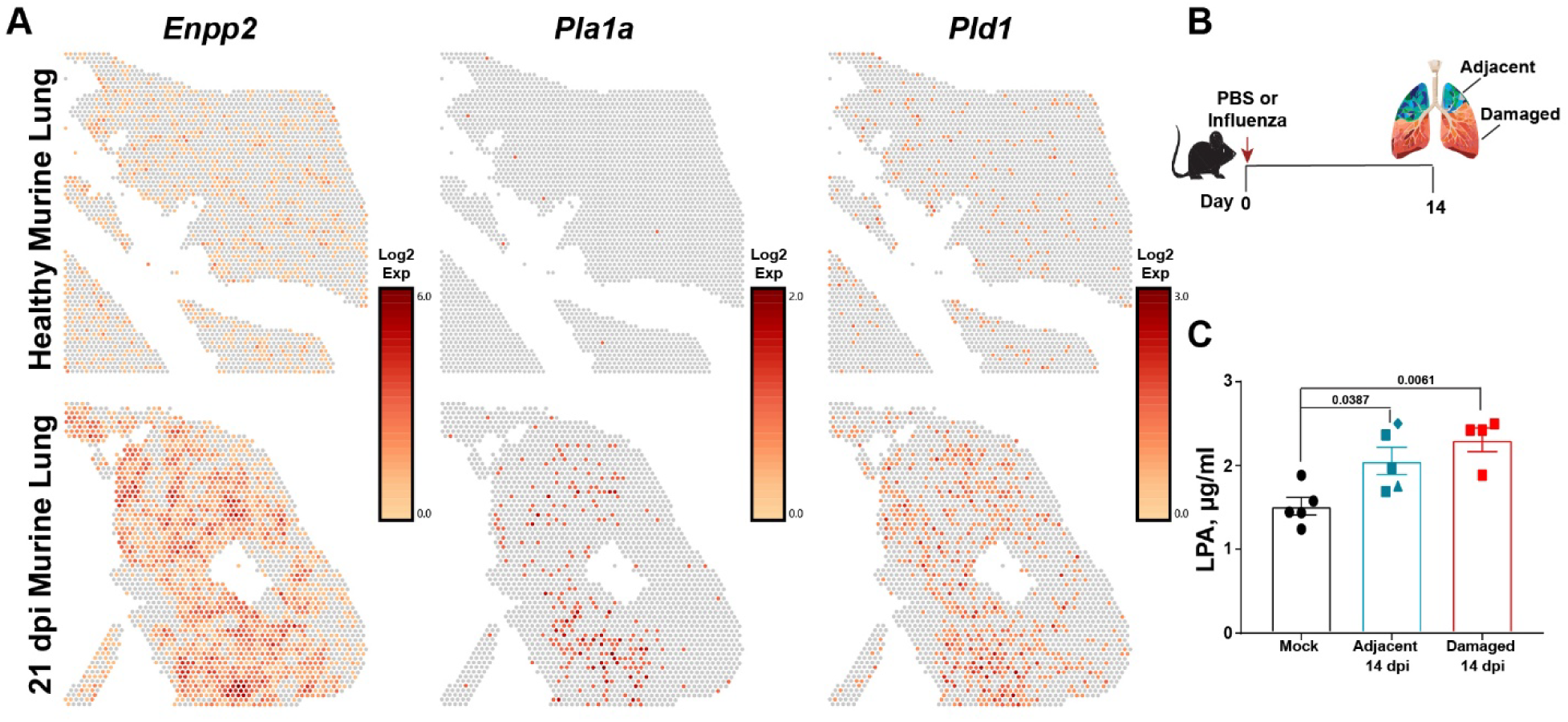
Influenza exposure induces LPA production and accumulation in damaged tissue. **(A)** Enzymes responsible for LPA production, such as Autotaxin (*Enpp2*), Phospholipase A1 (*Pla1a*) and Phospholipase D1 (*Pld1*) are highly expressed in post-influenza infected murine lung (dpi 21) as determined by 10x Visium Spatial Transcriptomics. Dataset was previously generated in our laboratory and is publicly available.^22^ Color intensity indicates the degree of log_2_ expression of the indicated genes. **(B)** Experimental design for quantification of LPA in tissue. C57BL/6 mice (7-11 weeks old, 7 male and 3 female) were infected with 12.5-15 units of Influenza virus. At 14 days post-flu (dpi), lung lobes were collected, homogenized and used for ELISA. Samples were combined from two independent experiments. R^2^ for standard concentration curves for both experiments is 0.98. **(C)** Bar plot shows increased LPA levels in damaged lungs at 14 dpi as measured by ELISA. Data shown as mean LPA concentration ± SEM. Each dot represents one biological replicate (n=4-5). A single high outlier (6.6 µg/ml in damaged group) was identified using a two-sided Grubbs’ test (p-value < 0.05) and removed. P values were calculated using a one-way ANOVA with post-hoc Turkey multiple comparison test.

### LPA induces AP-1 signaling via LPAR3 and LPAR4 receptors

We next focused on identifying the signaling hubs activated downstream of LPA responsible for its stark effects on gene expression. Considering strong induction of *Fosl1* (which is largely restricted to the transitional state) (**Figure 2F**) and predicted activation of *Jun* (**Figure 1G**) by LPA, as well as predicted engagement other transcription factors such as Sp1 and Egr1 (**Figure S3**) which are reported to cooperate with AP-1,^23–25^ we focused on determining how LPA stimulation results in rapid *Fosl1* upregulation. LPA signals through 6 known G-protein coupled receptors, but the relevant receptor in AT2s ultimately promoting upregulation of *Fosl1* is unknown. We first determined relative expression levels of each LPA receptor. In agreement with published literature and our single-cell RNA-sequencing data^26^ (**Figure 4A**), RT-qPCR revealed the virtual absence of LPAR1 expression in AT2 cells compared to lung fibroblasts, and LPAR5 was undetectable in both groups (**Figure S4**). By contrast, mRNA levels of LPAR3, LPAR4 and LPAR6 were significantly expressed in AT2s, so we turned to pharmacological agents to further investigate the roles of these receptors (**Figure 4B**).

**Figure 4.**
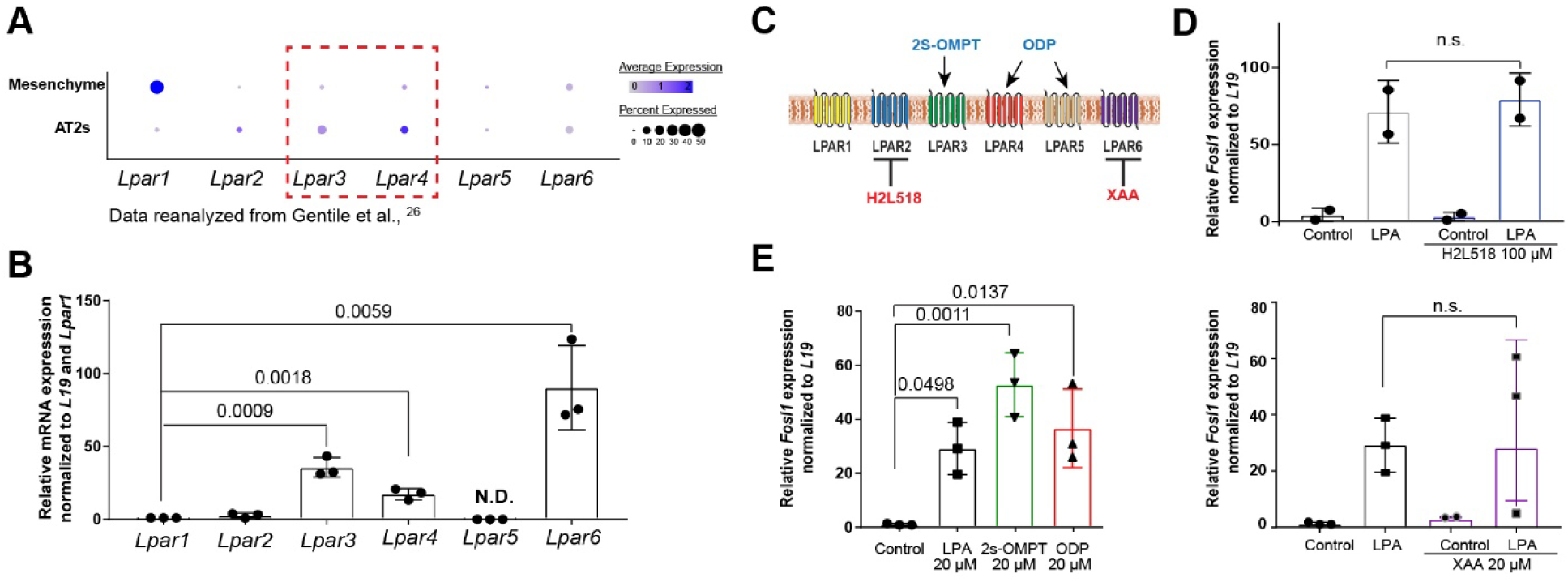
LPA induces AP-1 signaling via LPAR3 and LPAR4 receptors. **(A).** Dot plot of normal mouse lung non-immune cells for LPA receptors. Data reanalyzed from Gentile et al.^26^. **(B)** *Lpar1-6* expression was validated by RT-qPCR in 100k freshly sorted primary murine AT2 cells and normalized to *L19* and *Lpar1*. AT2s isolated as before from 11-12 weeks-old 2 female, 1 male mice. Data shown as mean fold change ± SEM (n.d. – not determined). Each dot represents one biological replicate (n=3). P values were calculated using a two-tailed unpaired t-test. **(C)** Schematic showing compounds that were used to selectively activate (blue) or inhibit (red) specific LPA receptors. **(D)** *Fosl1* expression as determined by RT-qPCR in murine AT2 organoids after treatment with LPAR agonists and antagonists, normalized to *L19*. At day 9 after seeding, AT2 organoids were starved for 22 hours, pretreated with 100 µM H2L5186303 or DMSO as a control and then were treated for 6 hours with 20 µM LPA in presence of 100 µM H2L5186303, ethanol treatment as a control. Data shown as mean fold change ± SEM. Each dot represents one biological replicate (n=2). P values were calculated using a one-way ANOVA with post-hoc Turkey multiple comparison test (n.s. – not significant). **(E)** *Fosl1* expression determined by RT-qPCR in murine AT2 organoids normalized to *L19*. AT2s were isolated by FACS from 3 C57BL/6 mice (13 weeks-old 2 female, 1 male) and plated on the top of matrigel (10000 cells per well). (*left*) At day 9 after seeding AT2 organoids were starved for 22 hours and followed by treatment for 6 hours with 20 µM LPA, 20 µM 2s-OMPT, 20 µM ODT or DMSO as a control. (*right*) At day 9 after seeding AT2 organoids were starved for 22 hours, pretreated with 20 µM 9-xanthenylacetic acid (“XAA”) and then were treated for 6 hours with 20 µM LPA in presence of 20 µM XAA, ethanol treatment as a control. Control and LPA samples presented on graphs (*left*) and (*right*) are identical as all experimental conditions were run simultaneously. Data show as mean fold change ± SEM. Each dot represents one biological replicate (n=3). P values were calculated using (*lef*t) a one-way ANOVA with post-hoc Turkey multiple comparison test and (*right*) two-tailed unpaired t-test (n.s. – not significant).

We treated murine AT2 organoids with selective LPARs agonists and inhibitors, with the effect of each drug presented in **Figure 4C**. Organoids were pretreated for 1 hour with each drug (or vehicle) followed by short-term LPA stimulation as in the earlier transcriptomic experiments. RNA collected from treated samples was used to perform RT-qPCR for *Fosl1* expression as our primary readout (**Figure 4D, E**). We observed that all tested agonists significantly upregulate *Fosl1* expression compared to vehicle control, where 2s-OMPT, an LPAR3 agonist, has a trend toward stronger induction (**Figure 4D (*left*)**). By contrast, we saw no effects of LPAR2 and LPAR6 inhibition tested by using pharmacological inhibitors H2L5186303 and XAA (**Figure 4D (*right*), E**). Based on these observations, LPA promotes AP-1 signaling in AT2s predominantly through LPAR3 and LPAR4.

### Impaired AP-1 signaling reduces AT2 proliferation

AP-1 is known to regulate myriad physiological processes including proliferation, apoptosis, differentiation, and response to hypoxia.^27,28^ We reasoned that the broad spectrum of phenotypes regulated by AP-1 support its potential role in controlling progenitor cell functions, which also involve diverse processes including proliferation, differentiation, and resistance to cell stress. However, the AP-1 family consists of 7 discrete transcription factors with overlapping and compensatory properties, making genetic deletion studies of individual AP-1 genes very difficult to achieve and interpret.^29^ Accordingly, to examine the necessity of AP-1 for effective alveolar regeneration by AT2s, we generated a novel transgenic mouse that is unable to activate all Jun-dependent AP-1 signaling due to inducible overexpression of a very well-validated dominant-negative form (**Figure 5A**) of c-Fos (AFOS). ^30–33^ We generated mice carrying this construct at the Rosa26 locus preceded by LoxP-STOP-LoxP (**Figure S5)**. To ensure the stability of the established mouse line, we isolated DNA after ≥ 4 generations of breeding, amplified the full-length AFOS construct, and confirmed 100% sequence identity with the original construct (**Figure S5**).

**Figure 5.**
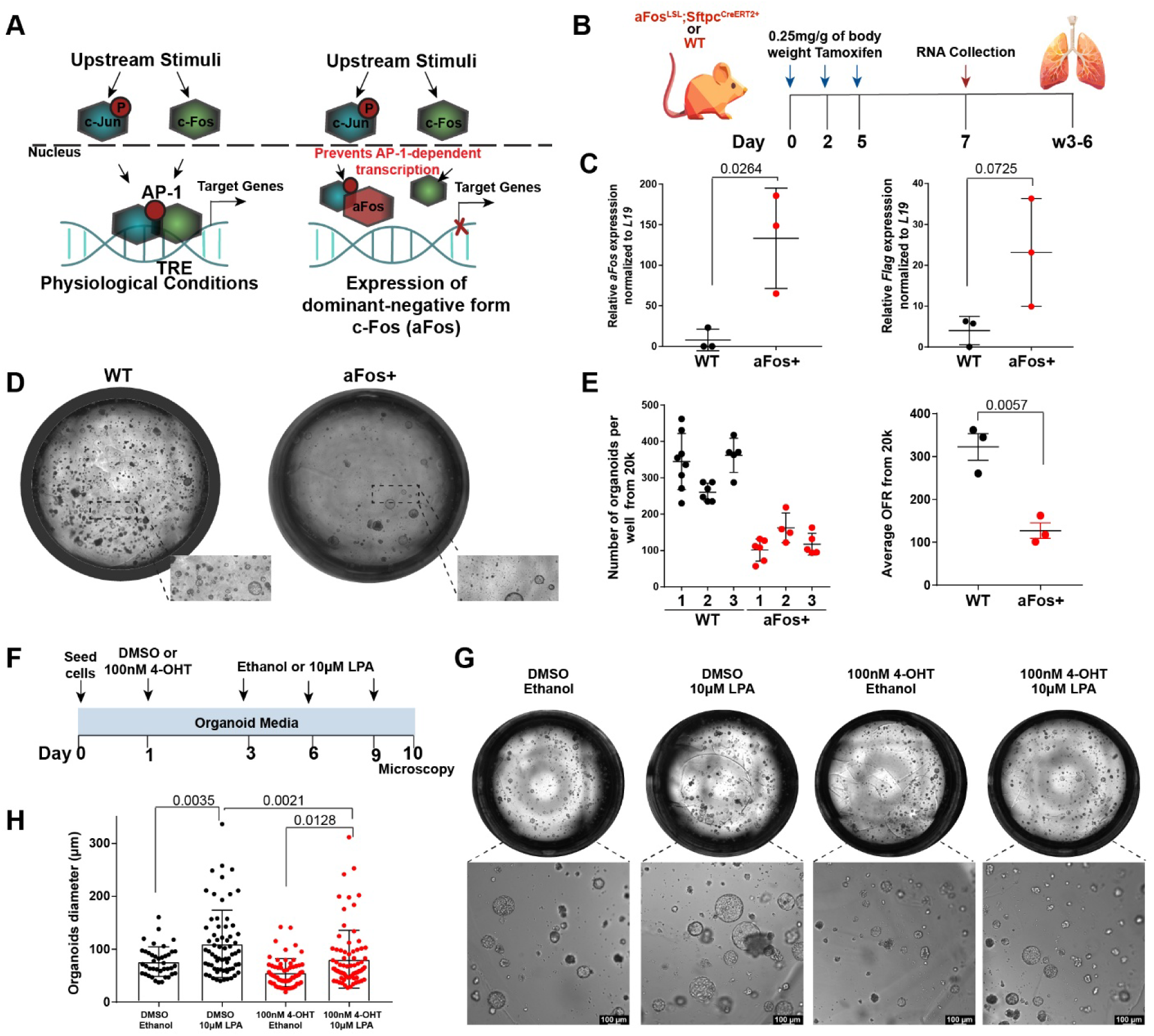
Expression of aFos construct in murine AT2 cells greatly diminishes progenitor function. **(A)** Schematic demonstrating mechanism of aFos action. Under normal conditions extracellular ligands can activate signaling cascades followed by phosphorylation of c-Jun (*left*). Phosphorylated c-Jun and c-Fos formed a heterodimer AP-1, which then binds TRE (TPA-response element) and activates expression of AP-1 target genes. Expression of a dominant-negative form of c-Fos (aFos) competitively binds to Jun and prevents AP-1-associated genes expression (*right*). **(B)** Schematic outline of the experimental design. aFos^+^; Sftpc^CreERT2+/+^ or functionally wild-type (wt, negative for at least one of the two transgenes) mice were treated with 0.25 mg/g of body weight tamoxifen by i.p. at days 0, 2 and 5. At day 7 after first injection, lung lobes were collected for RNA extraction. In addition, AT2s cells were isolated by FACS 3-6 weeks after tamoxifen administration to assess organoid formation. **(C)** Relative mRNA level of *aFos* and *Flag* normalized to *L19* in powder of lung lobes collected at day 7 from first i.p. injection. Each experimental group contained both male and female animals. Data shows mean fold change ± SEM. Each dot represents one biological replicate (n=3 mice). *P* values were calculated using an unpaired two-tailed t-test. **(D)** Microscopic evaluation of an entire well of organoids formed by AT2s derived from aFos+ or control (wt) animals, day 10. **(E)** Number of organoids formed from 20k cells represented as organoid formation rate at day 10-13 (OFR). Each dot represents OFR from one well, bar graph shows mean ± SD for one biological replicate (*left*). Bar graph shows mean ± SEM for average of OFR (*right*). Each biological represents one biological replicate. *P* values were calculated using an unpaired two-tailed t-test. **(F,G,H)** Freshly derived AT2 cells from aFos^+/+^; Sftpt^CreERT2+/+^ (aFos+) were plated on top of matrigel (15000 cells/well). Cre expression was initiated by treatment with 100nM 4-OHT or DMSO as a control on day 1. On day 3, media was replaced and supplemented with 10µM LPA or Ethanol as a vehicle control. Media was changed every other day. **(G)** Representative images of formed organoids. Microscopy imaging was performed at day 10 (x10). Scale bars, 100 µm. **(H)** Bar graphs represent average organoid diameter. Each dot represents one organoid (n=40-79 per condition). Data shown as mean ± SD (n=1). P values were calculated using a one-way ANOVA with post-hoc Turkey multiple comparison test.

To initiate *AFOS* expression only in AT2s, we crossed this mouse with tamoxifen-inducible Sftpc^CreERT2+^ (**Figure S5**) mice. Animals were treated with 3 doses of 0.25 mg/g of body weight tamoxifen or vehicle control by intraperitoneal injection (i.p.) and flash-frozen lungs were collected for analysis (**Figure 5B**). Inducible high expression of *AFOS* and its associated *Flag* tag were validated by RT-qPCR (**Figure 5C**). Notably, long term expression of AFOS in AT2s did not result in any obvious pathologic effects at homeostasis, with mice appearing overtly healthy through 6 months of AFOS expression (**Figure S5**).

To assess how AP-1 blockade impacts AT2 progenitor function *ex vivo*, we isolated primary AT2 cells from aFos^+^;Sftpc^CreERT2+^ and subjected these cells to organoid culture as described above. Blockade of AP-1 drastically reduced the organoid-forming capacity of AT2s (**Figure 5D,E**). In keeping with the predicted effects of AP-1 on stem cell activity, these findings confirm that AP-1 is necessary for organoid formation by AT2s.

Next, we asked whether the positive regulation of the organoid formation capacity of AT2s by LPA requires AP-1. We derived AT2s from aFos^+^;Sftpc^CreERT2+^ and established cultures for 24 hours. We then added 100nM 4-OHT or DMSO to induce recombination for 48 hours followed by treatment with 10 µM LPA or vehicle control through day 10 (**Figure 5F**). Blockade of AP-1 by *AFOS* largely, though not completely, prevented the trophic effects of LPA on AT2 organoid size (diameter) (**Figure 5G,H**). We conclude that AP-1 is a critical hub through which the pro-regenerative effects of LPA signal, though it remains possible that parallel pathways downstream of LPA also contribute to these effects.

### Impaired AP-1 signaling in AT2s prevents effective recovery from influenza injury

To address whether the absence of AP-1 signaling affects alveolar repair and the response to viral injury *in vivo*, we induced *AFOS* expression in aFos^+^;Sftpc^CreERT2+^ mice, allowed 21 days for *AFOS* induction and tamoxifen clearance, and infected with influenza (**Figure 6A**). AFOS expression in AT2 cells very significantly exacerbated body weight loss (**Figure 6B**) and decreased survival probability (**Figure 6C**) by 14 days post-infection, and AFOS-expressing mice still did not show full recovery even by 21 dpi (**Figure S6**). To further investigate the direct effects on AT2s, we isolated and quantified AT2s from AFOS and control mice at 14 dpi. We observed very reduced numbers of AT2 cells in AFOS+; Flu mice (**Figure 6D, Figure S7**) and high *AFOS* levels in the sorted AT2s (**Figure 6E**). These data indicate that genetic inhibition of AP-1 by AFOS expression prevents the normal regenerative properties of AT2s after viral injury.

**Figure 6.**
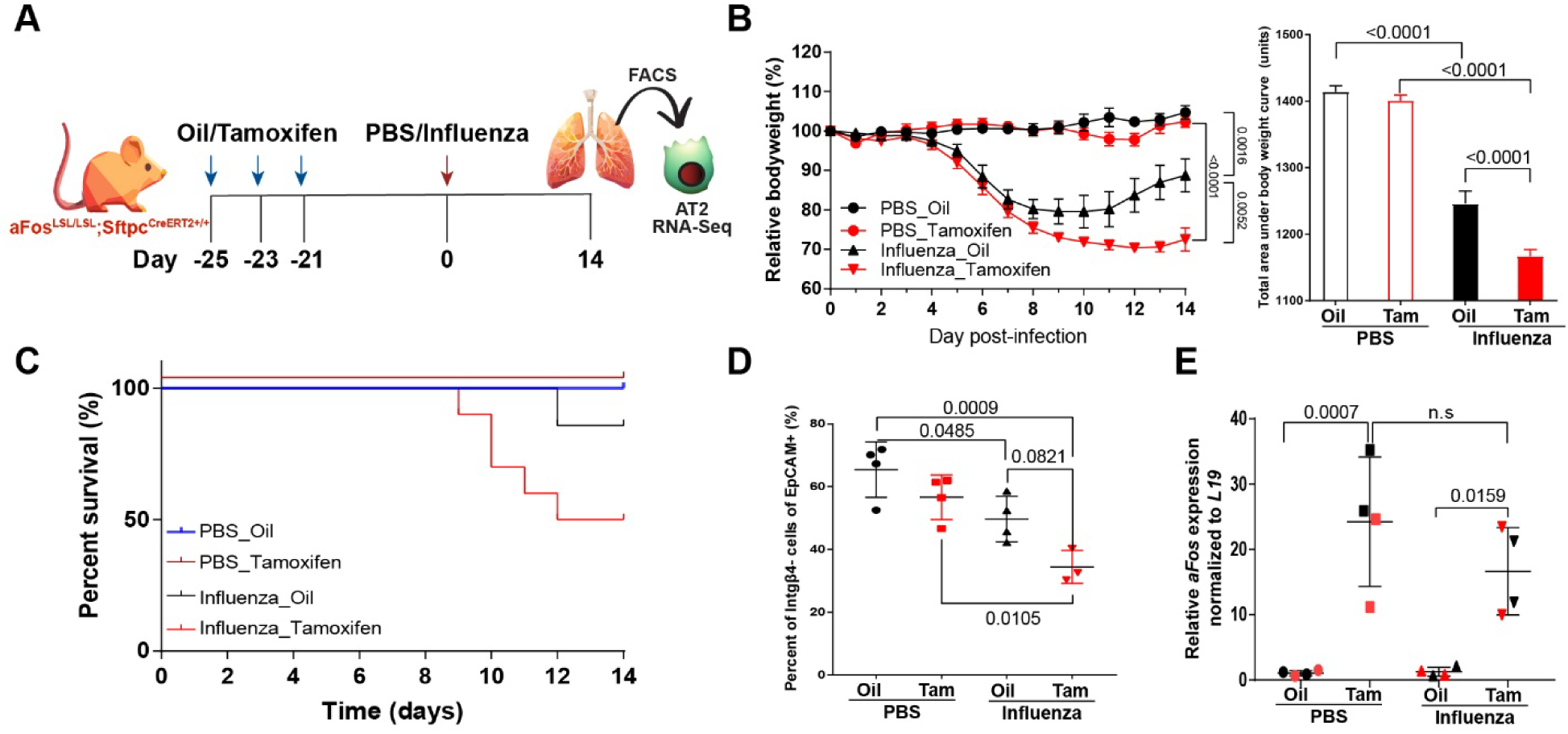
Genetic disruption of AP-1 signaling pathway in AT2s prevents normal recovery after influenza infection. **(A)** Schematic outline of the experimental design. For this experiment aFos^+/+^; Sftpt^CreERT2+/+^ (10F, 10M, 6-9 weeks old) mice and were equally divided between 4 groups: PBS_Oil, PBS_Tamoxifen, Influenza_Oil, Influenza_Tamoxifen. To induce expression of aFos, aFos^+/+^; Sftpt^CreERT2+/+^ mice were treated with 0.25 mg/g of body weight tamoxifen by i.p. or oil as a control at days −25, −23, −21. After 3 weeks, mice were infected with influenza virus. At 14 days post-flu (dpi), AT2 cells were isolated and sorted by FACS (draq7-; CD45-; EpCAM+; Integrin β4-) for further analysis from 4 mice per group (2F, 2M). **(B)** Body weight curve after infection. (*left*) Graph showing changes in relative body weight day normalized to day 0. Only mice that survived the entire experiment were included (n=4-7 per group). Values shown are mean ± SEM. P values were calculated for day 14 using a one-way ANOVA with post-hoc Turkey multiple comparison test. **(b)** Bar graph showing changes in relative units as area under curve, mean ± SD. P values were calculated using a one-way ANOVA with post-hoc Turkey multiple comparison test. **(C)** Kaplan-Meier survival curves for 4 experimental groups: PBS_Oil (blue), PBS_Tamoxifen (brown), Influenza_Oil (black), Influenza_Tamoxifen (red). **(D)** Scatter plot showing percentage of AT2s (Integrin-β4-) of the total EpCAM(+) epithelial population determined by flow cytometry. Data shown as mean ± SD. Each dot represents one biological replicate (n=4). P values were calculated using a one-way ANOVA with post-hoc Turkey multiple comparison test. **(E)** Relative mRNA level of *aFos* normalized to *L19* in sorted AT2 cells. Data shown as mean fold change ± SD. Each dot represents one biological replicate (n=4, 2F, 2M in each group). P values were calculated using a one-way ANOVA with post-hoc Turkey multiple comparison test.

To more carefully dissect *how* AP-1 blockade affects the regenerative phenotype of AT2s during lung injury, we performed RNA-Seq on these isolated AT2s (**Figure 6A**). Principal component analysis indicated no separation between WT and *AFOS* AT2s in uninjured mice, reinforcing a model wherein AP-1 is dispensable for the homeostatic properties of AT2s. However, significant differences emerged upon infection (**Figure 7A**), highlighting the importance of AP-1 signaling in response to stress and injury. Over 800 genes were significantly differentially expressed in *AFOS* AT2s after influenza (**Figure 7B**). Analysis of the DEGs enriched in WT post-flu AT2s show strong pathway enrichment of cell cycle / proliferation genes (**Figure 7C**), strongly supporting our prediction that AT2s utilize AP-1 to effectively engage proliferative pathways when switching on their progenitor functions. Intriguingly, and somewhat counter to our predictions, we observed significant enrichment of a few genes in the post-flu *AFOS* AT2s that are associated with the transitional cell state (*Lgals3, Sprr1a*). This suggests that some markers associated with the transitional state may represent general AT2 stress markers engaged by AP-1-independent mechanisms.

**Figure 7.**
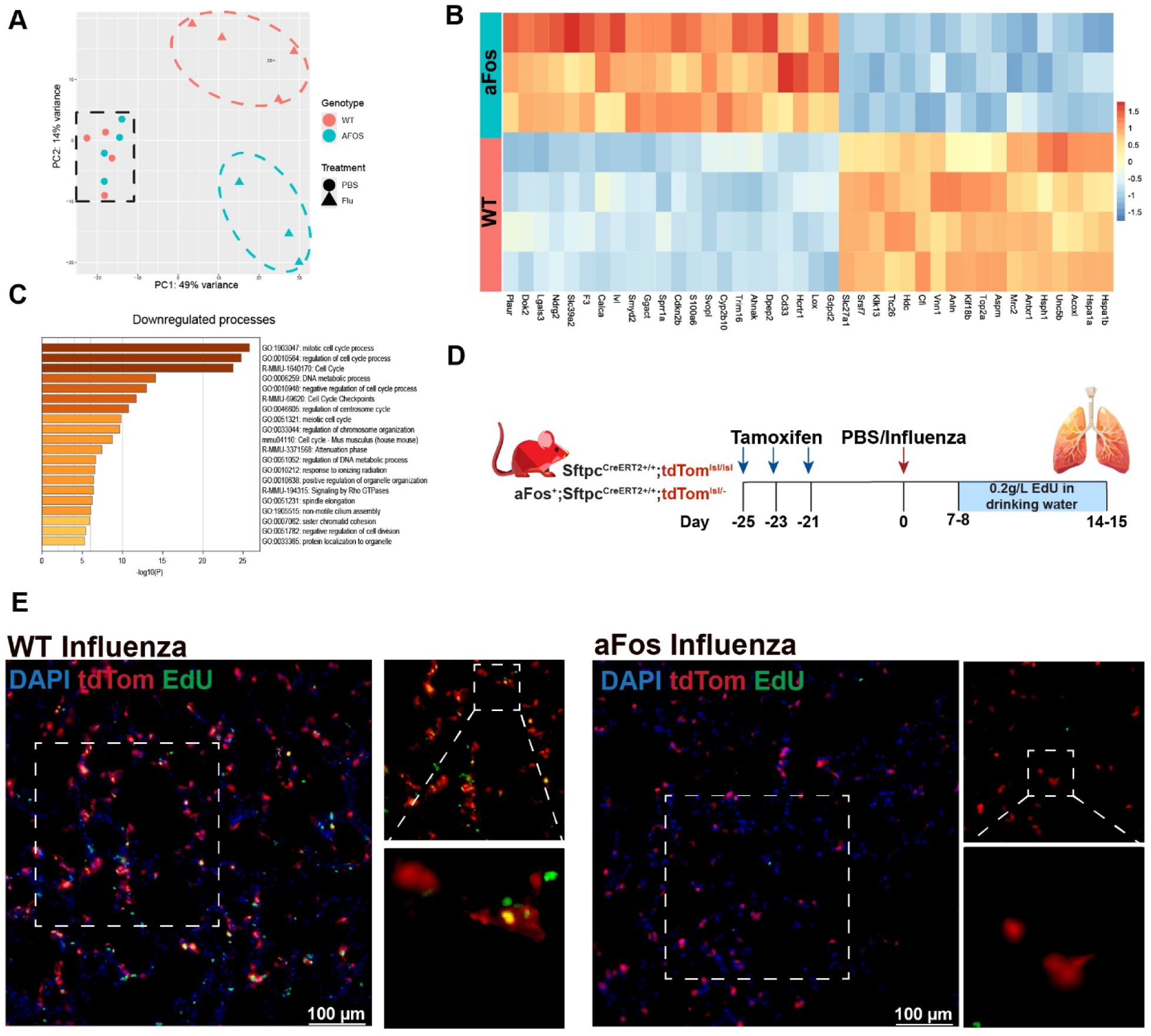
Genetic inhibition of AP-1 by aFos expression prevents normal recovery after influenza infection. **(A)** Principal component analysis (PCA) plot showing clustering for each sample used for bulk RNA-seq analysis. Clusters of samples based on infection status are outlined by dashed frame, where status on genotype was highlighted by color: orange for wild-type (WT) and blue for aFos+. **(B)** Functional enrichment analysis of the top significantly downregulated processes in aFos_Flu group compared to wild-type WT_Flu group determined via Metascape. **(C)** Heatmap for influenza-infected samples only. Top 40/882 DEGs (adj. p-value<0.05, LogFC>1). **(D)** Schematic outline of the lineage-trace experiment, including 4 groups: WT_PBS (3M, 3F; 6-12 weeks-old; mock-infected Sftpt^CreERT2+/+^; tdtomato^lsl/lsl^); WT_Flu (3M, 2F; 6-12 weeks-old; influenza-infected Sftpt^CreERT2+/+^; tdtomato^lsl/lsl^); aFos_PBS (2M, 4F; 6-12 weeks-old; mock-infected aFos^+^; Sftpt^CreERT2+/+^; tdtomato^lsl^); aFos_Flu (1M, 4F; 6-12 weeks-old; influenza-infected aFos^+^; Sftpt^CreERT2+/+^; tdtomato^lsl^). To induce expression of tdtomato and/or aFos, all mice were treated with 0.25 mg/g of body weight tamoxifen by i.p. at day −25, −23, −21. After 3 weeks, mice were infected with influenza virus. At 7-8 dpi, cage water was replaced with water containing 0.2g/L EdU. After 7 days (14-15 dpi) lungs were collected for histological analysis. **(E)** Representative immunofluorescence images from AT2 lineage-traced influenza-infected (Flu) mice that express aFos; tdtomato or tdtomato only. DAPI (Blue), EdU (Green), tdtomato (Red). Scale bars, 100 µm.

Lastly, we assessed the effects of AP-1 disruption on the proliferation capacity of AT2s *in vivo*. We incorporated a tdtomato^lsl/-^ allele into aFos^+^;Sftpc^CreERT2+^ mice to visualize AFOS+ AT2 cells and their progeny. aFos^+^;SftpcCreERT2+;tdtomato^lsl/-^ and SftpcCreERT2+;tdtomato^lsl/-^ (wt) mice were treated with 3 doses of 0.25 mg/g of body weight tamoxifen or vehicle and after 21 days from last injection were infected with influenza (**Figure 7D**). Mice had *ad libitum* access to water containing 0.2g/L EdU for 7 days prior to tissue collection to facilitate proliferation analysis. Lungs were collected at 14-15 dpi for histological evaluation and checked for presence of EdU incorporation by immunofluorescent microscopy. Our data revealed that influenza-infected AFOS+ mice have fewer AT2 (tdtomato+) cells per high-powered field compared to infected wt animals. Moreover, surviving AT2 cells in AFOS+ mice were less likely to have proliferated as judged by EdU incorporation (EdU+; tdtom+) (**Figure 7E**). This data confirm the transcriptomic analysis in Fig. 7C, indicating that AFOS expression *in vivo* impairs AT2s’ proliferation capacity. Taken together, this unique transgenic model implicates a central role for AP-1 signaling in the regenerative behavior of AT2s.

## Discussion

Here, we demonstrate that lysophosphatidic acid, a bioactive lipid generally associated with promoting fibrosis through its activity on fibroblasts, is sufficient to induce a regenerative state in the stem cells of the distal lung, alveolar type 2 cells. In contrast to dedicated stem cells in the skin or gut ^34,35^, AT2s exhibit replicative quiescence at homeostasis.^1,36^ As such, it follows that an external signal is needed to “wake up” the progenitor features of these cells. Our work here indicates that the distinct transcriptomic state generally referred to as the transitional state in fact represents exactly this scenario, in which the AT2 progenitor features have been switched on to allow for survival, proliferation, and differentiation in the response to alveolar injury (**Figure 8**). Further, we show that LPA is capable of switching cells into this state extremely rapidly, requiring no more than 6 hours of LPA stimulation to adopt an entirely distinct transcriptome, dwarfing the changes observed in response to other known signals implicated in the transitional state, namely IL-1β or p53.

**Figure 8.**
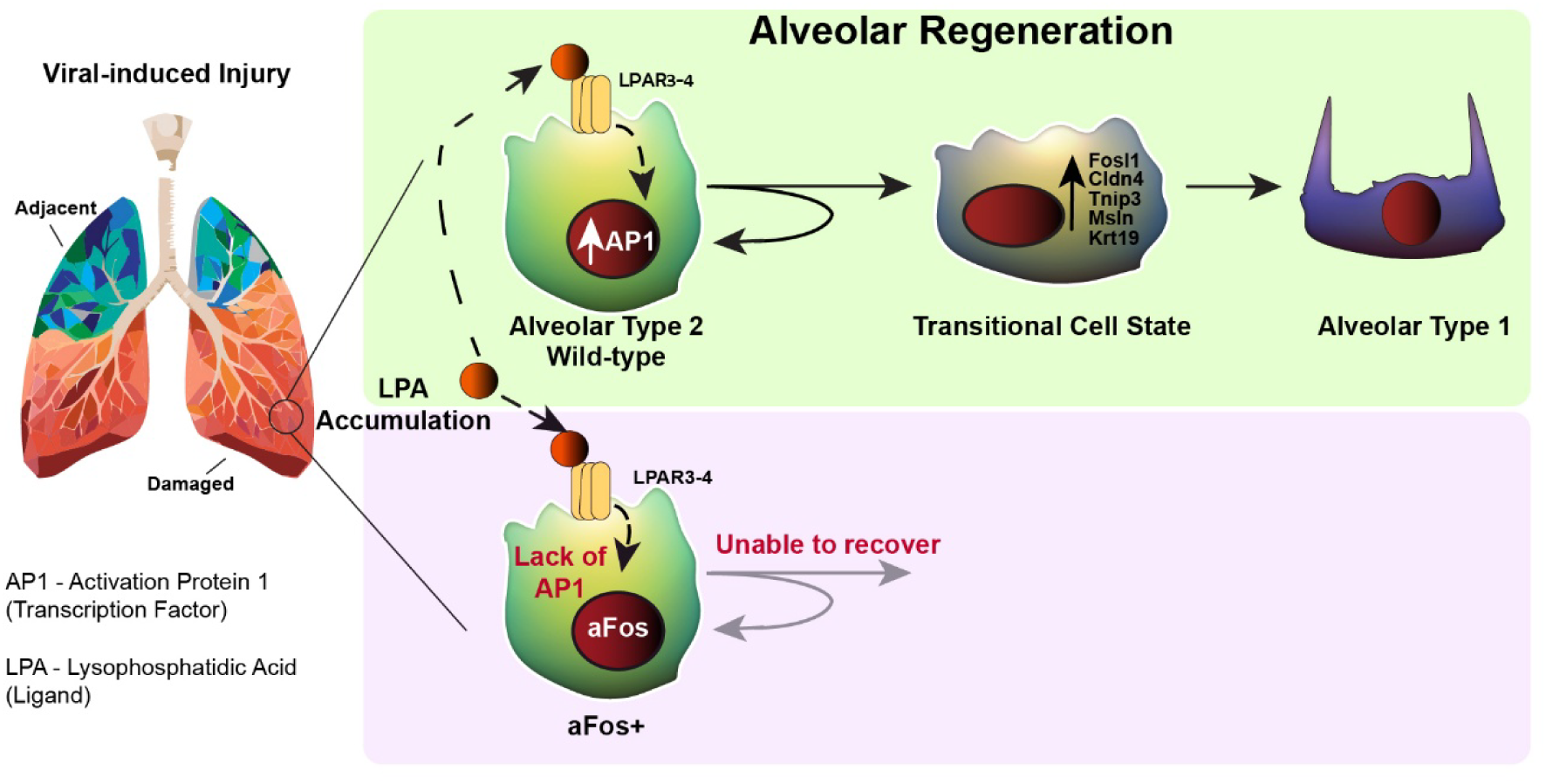
Proposed model

Downstream of LPA, our work points to a central requirement for AP-1 signaling in switching on AT2 progenitor features. The initial hypothesis suggesting a key role for AP-1 came from the observation that the core AP-1 family member *Fosl1* is the most highly upregulated transcriptional factor in AT2s after LPA stimulation, an observation further corroborated by *Fosl1* expression being essentially restricted to transitional AT2s *in vivo.* Given the diverse functions of AP-1 and the fact that AT2 cells in a progenitor state must likewise adopt vastly different functions than homeostatic AT2s (e.g. proliferation, differentiation, and motility vs serving as factories for surfactant production), AP-1 would indeed seem a prime candidate for controlling such progenitor features. Indeed, recent work from another group implicated AP-1 as being required at the epigenetic level for achieving entry into the transitional state, corroborating key aspects of our model. However, AP-1 factors are known to have tremendous redundancy with one another. Indeed, AT2-specific deletion of *Fosl1* did not result in any obvious phenotype upon influenza injury (**Figure S8**), and combined deletion of *Fos, FosB, and JunB* in AT2s in the aforementioned study resulted in a relatively mild cell dispersion phenotype after Sendai virus injury, with no reported changes in AT2 proliferation or animal morbidity / mortality. Conversely, by engineering a very well-studied and validated dominant negative AP-1 construct into AT2 cells, ^30–33^ we observed a striking regenerative failure manifest at both the cellular (reduced proliferation and survival) and organismal (increased mortality and failure to recover body weight), validating the idea that AP-1 mediated gene expression is absolutely critical for adaptive responses to alveolar injury.

Though widely observed and conserved in many injury models, the fundamental nature of the AT2 transitional state remains highly debated. Arguments for this state being largely pathological revolve around observations of these cells being initially observed in fibrotic injury models (e.g. bleomycin),^37,38^ evidence of a senescence signature / p53 activation (which would be expected to slow or halt proliferation)^8^, and their observed secretion of proinflammatory or profibrotic mediators that might be expected to promote pathology.^39^ However, the idea that that this state is inherently pathologic needs to be qualified by the lenses in which these observations were made. First, temporary promotion of fibrotic processes is part of normal wound healing, e.g. deposition of provisional matrix, and should only be viewed as pathologic if it is persistent or uncontrolled. In keeping with this, several studies observing the transitional state purposefully drive expression of pathologic gene variants without the ability to downregulate these toxic gene products, thus locking cells into a terminally stressed condition.^40–44^ Finally, our *in vivo* transcriptomic data with the dominant negative AP-1 indicates that some genes associated with the transitional cell state (*Lgals3*, *Sprr1a*) can still be upregulated in the absence of AP-1-medaited transcription, so it may be more appropriate to consider these genes general AT2 cell stress indicators more than markers of a transitional progenitor state.

It is intriguing that a molecule so classically associated with promoting pulmonary fibrosis, LPA, has such strong pro-regenerative effects on AT2 cells. Our findings suggest that like the transitional state itself, instead of being inherently pro-pathologic, LPA should be considered a key signal in the normal reparative process which becomes pathologic only in excess. Our model suggests the ability to “tune” the distinct responses to LPA in epithelial vs mesenchymal progenitors has probably been selected for evolutionarily, resulting in six distinct LPA receptors which are by and large mutually exclusive between fibroblasts and AT2s. Indeed, our data indicating the pro-regenerative effects of LPA on AT2s operates largely through LPAR3 and LPAR4 but not LPAR1, which is restricted to fibroblasts, highlight an exciting opportunity to utilize or design receptor-specific ligands that will promote AT2 regenerative behavior without activating fibrotic responses.

### Study Limitations

Like any new work, our studies are not without their limitations. We utilize an H1N1 viral pneumonia model throughout this work, and it is formally possible that the effects of AP-1 and LPA could differ in response to different injuries. Though we validated upregulation of *Fosl1* in response to LPA in human AT2s, we cannot otherwise directly assess the role of AP-1 in human lung injury *in vivo*. That being said, another group has similarly implicated AP-1 in adoption of the human “basaloid” state generally thought to be synonymous with or at least related to transitional AT2s in mice, broadly supporting the evolutionary conservation of our findings.^29,45^ Further, it is important to note that we do not claim that LPA is the *only* signal capable of promoting AT2 progenitor cell activation, just that it is a very robust signal that appears to be sufficient do so in isolation, given that short term stimulation with LPA experiments were performed after starvation with a basal ADMEM/F12 media. Given the diversity of paracrine signals present in injured lungs,^4^ it is of course possible that additional ligands cooperate with LPA to promote AT2 progenitor activation, and that the role of discrete signals may be specific to the nature of the injury.

In sum, our work indicates a central role for both lysophosphatidic acid and AP-1 in switching on the progenitor features of alveolar type 2 cells. Further, the biological serendipity of differential receptor expression and responsiveness between fibroblasts and AT2s highlights an exciting opportunity through which therapeutic engagement of epithelial regeneration might be possible without accompanying fibrotic activation of mesenchyme. Future studies should focus on cell type-specific LPA stimulation and potential disentanglement of various components of AP-1 transcriptional regulation to further promote euplastic alveolar re-epithelization while limiting fibrotic outcomes of severe lung injury.

## Methods

### STAR Methods

#### Mice

All murine studies were performed under a University of Pennsylvania Institutional Animal Care & Use Committee-approved protocol (#806262). Mice were housed in individually ventilated caging racks on corncob bedding with 12:12 light-dark cycle housed in an Animal Biosafety Level 1 (ABSL-1) facility. Wild-type C57Black/6J (C57BL/6 in this manuscript) (Cat# 000664) mice were purchased from Jackson Laboratories. *aFos^LSL^* mice were generated in our laboratory. *aFos^LSL^* mice were crossed with Sfptc^CreERT2+/+ 46^ and *Rosa26R*^lsl-Ai14-tdTomato(tdTomato) 47^. The primers used for genotyping are listed in **Table S2**. DNA was isolated from tail clips using tissue preparation solution (Sigma-Aldrich, Cat# T3073), extraction solution (Sigma-Aldrich, Cat# E7526), neutralization solution (Sigma-Aldrich, Cat# N3910) following manufacturer’s protocol. Notably, since both *aFos* and *tdTomato* transgenes were inserted in the same Rosa26R locus, interpretation of genotyping results is slightly different than usual. Mice cannot be homozygous for both genes at the same time.

### Generation of *aFos^LSL^* (*Rosa26R*^lsl-aFos^) mice

*Rosa26R*^lsl-aFos^ (*aFos^LSL^*) mice were generated in V6.5 mouse ESCs as described earlier.^48^ Mouse aFos cDNA was amplified from CMV500 A-FOS plasmid (Addgene, #33353) by PCR. aFos PCR products were cut, purified, and inserted into a generic targeting vector (pBigT, Addgene, #21270), pBigT-aFos was then subsequently cloned into a plasmid with the ROSA26 genomic flanking arms (ROSA26-PA, Addgene, #21271) following the protocol described previously,^49^ generating the final targeting vector ROSA-26-pBigT-aFos for homologous recombination. The linearized targeting vector was electroporated into mouse ESCs (same line as described above), and selected colonies were analyzed for proper editing by PCR. Finally, the targeted ESCs were injected into C57BL/6J blastocysts. Chimeric mice (*aFos^LSL^*) were bred with Sfptc^CreERT2+/+^ male mice for further characterization. Genotyping primers and PCR program are listed in **Table S2**. To ensure that established mouse line did not mutate over time, we isolated DNA from a murine tail after greater than 4 generations of breeding, we amplified the full-length aFos construct using forward primer: TGGACTACAAGGACGACGAT; reverse primer: ACAGTCGAGGCTGATCAGCGA. Sequence of this PCR product (293 bp >100 ng) was performed by Plasmidsaurus using Oxford Nanopore Technology with custom analysis and annotation and compared with original plasmid (CMV500 A-FOS, Addgene, #33353) by using BLAST tool.^50^ Identity is 100% (293/293), **Figure S3**.

### Tamoxifen-treatment and Influenza lung injury

A minimum of three animals per group was used for studies involving statistical analyses. Sample size was determined by availability and previous experience with influenza infection experiments in mice. The current study includes several transgenic mouse models: *aFos^LSL^*; Sfptc^CreERT2^ (aFos+ AT2 cells), Sfptc^CreERT2^ (WT control for aFos+ mice), *aFos^LSL^*; Sfptc^CreERT2^; *Rosa26R*^lsl-Ai14-tdTomato(tdTomato)^ (lineage-tracing for aFos+ AT2 cells), Sfptc^CreERT2^; *Rosa26R*^lsl-Ai14-tdTomato(tdTomato)^ (AT2 lineage-trace control). Exact genotype, age, sex, number of mice are provided in corresponding figures and figure legends. At least 6-week old mice were administered three doses of tamoxifen (250 mg kg^−1^ body weight) in 50 μl of corn oil every other day and rested for 3 weeks after the last injection. Then mice were transferred to the Animal Biosafety Level 2 facility at the University of Pennsylvania. Afterward, influenza virus A/H1N1/PR/8 was administered intranasally at 12.5–17.5 TCID_50_ units of median tissue culture infectious dose to mice (15-20 g, 12.5 U; 20–25 g, 15 U; 25–30 g, 17.5 U) dissolved in 30 μl of PBS as we previously described.^51,52^ Control mice were administered 30 μl of PBS. Mice were weighed regularly. Diet gel (Clearh2o, #72-06-5022) was provided to all animals when at least 1 mouse in the ongoing experiment lost 15% of body weight. Mice that lost 15% or more of their initial body weight by day 9 were deemed adequately infected and used for all experiments involving influenza infection. Animals were euthanized and lung tissues were collected at the specified timepoints (majority at 14 or 21 dpi) for further analyses including histology, FACS, ELISA or RT-qPCR.

### Immunofluorescent lineage tracing and 5-ethynyl-2’-deoxyuridine (EdU) treatment

To initiate labeling of AT2 cells, *aFos^LSL^*; Sfptc^CreERT2^; *Rosa26R*^lsl-Ai14-tdTomato(tdTomato)^ and Sfptc^CreERT2^; *Rosa26R*^lsl-Ai14-tdTomato(tdTomato)^ mice were administered three doses of tamoxifen (250 mg kg^−1^ body weight) in 50 μl of corn oil every other day and rested for 3 weeks after the last injection, resulting in lineage-traced AT2s. Mice were then infected with influenza as described above or PBS as a mock control. For detection of cell proliferation *in vivo*, 0.2g/L of EdU (GoldBio, #E-980-1) was dissolved in sterile drinking water. Mice had *ad libitum* access to filter-sterilized water for 7 days starting at 7-8 dpi. Water has been changed every 2-3 days. This experiment included 4 groups: WT_PBS (3M, 3F; 6-12 weeks-old; mock-infected Sftpt^CreERT2+/+^; tdtomato^lsl/lsl^); WT_Flu (3M, 2F; 6-12 weeks-old; influenza-infected Sftpt^CreERT2+/+^; tdtomato^lsl/lsl^); aFos_PBS (2M, 4F; 6-12 weeks-old; mock-infected aFos^+^; Sftpt^CreERT2+/+^; tdtomato^lsl^); aFos_Flu (1M, 4F; 6-12 weeks-old; influenza-infected aFos^+^; Sftpt^CreERT2+/+^; tdtomato^lsl^). Lungs were collected at 14-15 dpi for histology as described in the following section. Fixed frozen sections of lungs were stained with Click-iT EdU Imaging kit with Alexa Fluor 488 Azide (Thermo Fisher Scientific, #C10086) according to the manufacturer protocol. EdU+ cells were detected by fluorescent microscopy.

### Histology and Immunostaining

After euthanasia with approved methods, the lungs were thoroughly perfused with cold PBS via the left atrium to remove residual blood in the vasculature, inflated with 3.2% PFA for 1 min and then with 2% UltraPure Low Melting Point Agarose (Invitrogen, #16520050) in PBS for 2 min. Lungs were further fixed in 3.2% PFA for 1 h at room temperature on a shaker (100 rpm), rinsed three times with PBS for 20 min each and incubated overnight at 4°C in 30% Sucrose in PBS (Millipore Sigma, #S8501) with 0.02% sodium azide (Millipore Sigma, #S2002). Lungs were then transferred to 15% sucrose 50% optimal cutting temperature (OCT, Thermo Fisher Scientific, #23-730-571) compound and incubated at room temperature for 3 h. Fixed tissues were transferred to an embedding mold filled with OCT compound, flash frozen in a dry ice ethanol bath, and stored at −80°C. Frozen sections (7 µm thick) were cut by Leica CM3050 S Research Cryostat (Leica Biosystems) at −20°C, and slides were stored at −20°C until being used for immunostaining as described earlier.^14^ Briefly, tissue sections were warmed to RT for 10 min and further fixed for 5 min in 3.2% PFA, rinsed two times with PBS and incubated in PBS for 30 min at 55 °C to remove residual agarose. Slides were then incubated with 0.5% triton X-100 (Fisher Bioreagent, Cat# BP151-100) in PBS for 10 min and washed three times with filtered 3% BSA (GoldBio, Cat# A-429-100) in PBS. 100 µl of reaction mix was added to each section, incubated for 45 min at dark at 37°C on a shaker at 20 rpm. After washing three times with filtered 3% BSA, slides incubated in 1 μM DAPI (Fisher Scientific, D1306) for 5 minutes, rinsed with PBS, and mounted with Fluoroshield (Sigma Aldrich, Cat# F6182), dried overnight at room temperature and store at 4°C until imaging.

### Imaging and Quantification

Images were taken on a Leica DMi8 microscope with a Leica DFC9000 sCMOS camera using Leica Application Suite X (LAS X) software. Immunofluorescent and phase contrast images were taken at 10x and 20x. Immunofluorescence microscopy was performed for selected regions (3-5 high-powered fields per lobe) to determine presence of tdtomato-expressing and EdU+ cells.

Phase contrast images of organoids were taken at 10x and 20x. Number of organoids per well and organoid diameters were measured by FIJI (ImageJ, 1.54p). For **Figure 2J**, microscopic imaging was performed at day 8 after seeding. Five high-powered fields were analyzed for each condition. Average diameter of at least 29 organoids per each biological replicate was used for statistical analysis (n=4; 2M, 2F). For **Figure 5D**, microscopic imaging was performed at day 10-13 after seeding. Number of organoids formed from 20k cells in each well was calculated. For **Figure 5G**, microscopy imaging was performed at day 10 (x10). The diameter of at least 40 organoids was measured per condition to define the average.

### ELISA

Influenza-infected wild type C57Bl/6J mice (or uninfected control mice) were euthanized at 14 dpi. The lungs were thoroughly perfused with cold PBS via the left atrium to remove residual blood in the vasculature. Lung lobes were separated, transferred to an eppendorf tube and centrifuged at 1,000 × g for 1 min at RT to remove remaining PBS. Then we manually separated and cut off by razor damaged and adjacent areas and weighed them separately. Cold PBS was added to each sample to achieve 0.6 mg/µl of tissue and was carefully homogenized on ice 5 times with an electronic homogenizer (L-HOB-Mini, Labgic). Material was centrifuged at 1,500 × g for 15 min at 4°C. Collected supernatant was stored at −80 until use. A commercially available ELISA kit (MyBioSource, Cat# MBS7269921, lot 20240701) was used to quantify mouse lysophosphatidic acid (LPA). All reagents, samples and standards were prepared according to the manufacturers’ instructions and 100 µl of samples were added to a well in pre-coated 96-well plates provided with the kit. The contents of the plates were then measured at a wavelength of 450 nm using the GloMax Discover plate reader (Promega). Data from each plate were calculated using standard curve run in each plate (R^2^=0.98). Each group has n=5, however one sample from “damaged” group was excluded due to high concentration verified by outlier test (p<0.05).

### Whole lung single cell suspension preparation, AT2 isolation by fluorescence-activated cell sorting (FACS)

#### Murine AT2s

Murine AT2 isolation methodology was performed as previously described, with minor modifications described here.^14^ Lung cells were isolated by first inflating lungs PBS and then with 15 U/mL dispase II (Gibco, Cat#17105-041) in HBSS (Gibco, Cat#14175-079), tying off the trachea, and cutting lobes away from the main stem bronchi. Lobes were then incubated in dispase for 45 minutes shaking at room temperature and mechanically dissociated by pipetting in FACS buffer (phenol-free Dulbecco’s modified Eagle’s medium (DMEM, Gibco, Cat# 21063-029); + 2% cosmic calf serum (Thermo Fisher Scientific, Cat# SH30087.04HI) + 1% penicillin streptomycin (p/s, Gibco, #15140-122). After pelleting at 550 × *g* for 5 minutes at 4°C, whole-lung suspension was treated with Red Blood Cell (ACK) Lysis Buffer (Gibco, Cat# A10492-01) for 4 minutes on a shaker at 37°C, washed with 10 ml FACS buffer and pelleted. Cell count was performed manually under microscope using Trypan Blue solution (1:1 dilution) to evaluate percentage of dead cells. For all reported experiments in this study, percentage of live cells was ≥95%. In order to isolate murine AT2 cells by FACS, whole lung single-cell suspensions from (a) *aFos^LSL^*; Sfptc^CreERT2+^; (b) *aFos^LSL^*; (c) Sfptc^CreERT2+^ or (d) C57Bl/6J mice were blocked in PBS containing 1:50 mouse TruStain FcX (anti-mouse CD16/32, BioLegend, Cat# 101320, clone 93) for 10 min on ice. The cell suspension was stained for allophycocyanin (APC)/Cy7-conjugated rat anti-mouse CD45 antibody (1:200, BioLegend, 30-F11, Cat# 103116, lot B452900), Brilliant Violet rat anti-mouse CD326 (EpCAM) antibody (1:100, BioLegend, G8.8, Cat# 118225, lot B424488), phycoerythrin (PE)-conjugated rat anti-mouse CD104 (Integrin-β4) antibody (1:100, BioLegend, 346-11A, Cat# 123610, lot B379262), and 10 nM of LysoTracker green DND-26 (Invitrogen, Cat# L7526, lot 2272635). Staining was performed for 30 minutes at 4 °C, followed by two washes with FACS buffer and a final spin at 550 × *g* for 5 minutes at 4 °C. Stained cells were then resuspended in FACS buffer containing DNAse (1:200 of 8900 U/ml, Millipore, Cat# D4627) and Draq7 (1:1000, BioLegend Cat# 424001, lot 304DR71000) as a live/dead stain. Each experiment included compensation and unstained controls. FACS sorting was done on Aria II (BD Biosciences) and cells were collected in Falcon™ round-bottom polystyrene tubes (Falcon Corning, Cat# 352054) into 1 ml of FACS buffer. Representative gating strategy is presented in **Figure S1**. Isolated cells were used for organoid formation assay and RNA isolation.

### Human AT2s

Human AT2s were derived from heathy donor’s lobe as described previously.^53^ The distal lung tissue was dissected into roughly 4 g pieces; the pleura and airways was removed by cutting and pulling with scissors and forceps. Tissue was washed with sterile PBS several times. After removing as much liquid as possible, the lung chunks was minced with scissors and Razor Blades into small pieces and transferred into four 50 ml tubes containing 10 ml of digestion buffer (540 U/ml Collagenase type 1, 5 U/ml Dispase,50 U/ml DNase I, 1X antibiotic-antimycotic (Gibco, Cat# 15240062) in HBSS) and digested for 2 h at 37 °C in a shaker at 300 rpm. Tissue was liquefied in the digestion solution using an Oster Blender as follows: (low setting for all) 5 s milkshake, 3 s smoothie, and 5 s milkshake. The digest suspension was poured through a funnel lined with sterile 4 × 4 gauze; filtered through a 70 µm strainer and spined down at 500 x g for 5-10 minutes. Next, 2ml of ACK lysing buffer was added to the cell pellet and mix gently for 2 minutes at room temperature and washed with 10 ml of wash buffer (0.5% BSA, 2mM EDTA in PBS), passed through a 40 µm cell strainer, and centrifuge at 500 x g for 5-10 minutes at room temperature. Cells were resuspended in 10 million cells/ml in freezing media: 60% FBS, 10% DMSO, 30% DMEM; slow froze down and stored in −80 °C. Prior to sorting cryovial was thawed in a 37 °C water bath. Then cells were resuspended in 10 mL 10% FBS (in DMEM); spun down at 500 x g for 5 min, and resuspended in 1 ml of wash buffer. Cell suspension was filtered through a 35 μm cell strainer, spun down and incubated with FcBlock (2.5 µl/10^6 cells/100µl; anti-FcR, BioLegend, Cat# 422302) for 10 minutes at room temperature. Next, cells were stained for 30 min on ice with following antibodies: Alexa Fluor anti-human CD326 (EpCAM) antibody (50 µg/ml, BioLegend, Cat# 324212, clone 9C4); PerCP/Cyanine5.5 anti-human CD45 antibody (50 µg/ml, BioLegend, Cat# 304027, clone HI30); primary anti-HT2-280 (10.26 µg/ml, Terrace Biotech, Cat# TB-27AHT2-280, Mouse IgM); secondary Alexa Fluor 488 anti-mouse IgM (500 µg/ml, BioLegend, Cat# 406522, clone RMM-1) and Draq7 (1:1000, BioLegend Cat# 424001, lot 304DR71000) as a live/dead stain. Stained cells were isolated by FACS using CytoFLEX SRT (Beckman Coulter) sorter. Representative gating strategy is presented in **Figure S2**. Isolated human AT2s (CD45-, Draq7-, EpCAM+, HT2-280+) cells were used for organoid formation assays.

### 3D-cell culture

FACS-sorted murine and human AT2 cells were seeded on top of 80 µl Matrigel (Corning, Cat# 354230) in “regular organoid media” containing: PneumaCult Alveolar Organoid Expansion Media (Stemcell, Cat# 100-0848), 10x supplement (1:10, Stemcell, Cat# 100-0849), 0.0005% heparin (Stemcell, Cat# 07980) and 1x penicillin / streptomycin. At initial seeding regular media was supplemented with “passaging compound” (1:100, Stemcell, Cat# 100-0860) only at initial plating. Later, media has been changed every 3 days with regular organoid media. For **Figure 2G**, human AT2 cells at day 9 after seeding AT2 organoids were starved for 22 hours with Advanced DMEM/F12 (“ADMEM/F12”, Gibco, #12634-010) and followed by treatment for 6 hours with 20 µM 1-Olyeyl-LPA (“LPA”, Cayman, Cat#10010093) or ethanol as a control. For **Figure 4E,D** freshly sorted AT2s were isolated by FACS from 3 C57BL/6 mice (13 weeks-old 2 female, 1 male) and plated on the top of matrigel (10000 cells per well). At day 9 after seeding AT2 organoids were starved for 22 hours and followed by treatment for 6 hours with 20 µM 1-Olyeyl-LPA (“LPA”, Cayman, Cat#10010093), 20 µM 2s-OMPT (Caynam Chemical, Cat# 1005707), 20 µM 9Z-Octadecenyl phosphate ammonium salt (“ODT”, Echelon Biosciences, Cat# L-0418) or DMSO as a control. To test the effect of LPARs inhibitors, at day 9 after seeding AT2 organoids were starved for 22 hours, pretreated with 20 µM 9-xanthenylacetic acid (“XAA”, Santa Cruz, Cat# sc-326122) and then were treated for 6 hours with 20 µM LPA in presence of 20 µM XAA, ethanol treatment as a control. Again, at day 9 after seeding, AT2 organoids were starved for 22 hours, pretreated with 100 µM H2L5186303 (Medchem express, Cat# HY107616) or DMSO as a control and then were treated for 6 hours with 20 µM LPA in presence of 100 µM H2L5186303, ethanol treatment as a control. For **Figure 5F-H**, freshly derived AT2 cells from aFos^+/+^; Sftpt^CreERT2+/+^ mouse were plated on top of matrigel (15000 cells/well). Cre expression was initiated by treatment with 100 nM 4-OHT ((E/Z)-4-hydroxytamoxifen, Cayman Chemical, Cat# 17308) or DMSO as a control on day 1. On day 3, media was replaced and supplemented with 10µM LPA or Ethanol as a vehicle control. Media was changed every other day. Microscopy imaging was performed at day 10. For other treatment conditions, including timeline and drug concentrations, relevant information is specified in the corresponding chapter “collection of samples and analysis for bulk RNA-sequencing” and in corresponding figure legends.

### RNA isolation and Quantitative qPCR

Total RNA from 15 mg flash frozen murine lung lobes was extracted by Direct-zol RNA Miniprep Plus kit (ZymoResearch, R2072) according to the manufacturer’s protocol. Total RNA from cultured cells, freshly isolated murine AT2 cells or organoids was extracted using the ReliaPrep™ RNA Cell Miniprep kit (Promega, #Z6011). The amount of RNA input for complementary DNA synthesis was standardized within each experiment to the RNA isolate with the lowest concentration per sample group as measured by Nanodrop (Thermo Fisher Scientific). cDNA was synthesized using the iScript Reverse Transcription Supermix (BioRad, #1708841) for **Figure 2G**, **4D**,**E** and **5C**, and by Takara SMART-Seq mRNA HT Kit (Takara, #634795) for **Figure 2F**, **4B**, **6E**. Quantitative qPCR was performed using PowerUp SYBR Green Master Mix (Applied Biosystem, #A25742) and ran with standard protocol on an Applied Biosystems QuantStudio 6 Real-Time PCR System (Thermo Fisher Scientific). Gene expression level was analyzed by the 2^-ΔΔCt^ comparative method and normalized to *Rpl19* (“L19”) or *GAPDH* for human. The used primers are listed in **Table S3**.

### Collection of samples and analysis for Bulk RNA-sequencing

#### 1. Bulk RNA-seq of organoids treated with Nutlin-3a, IL-1β and LPA

CD45-, draq7-, EpCAM+, Integrin-β4-, Lysotracker green+ single cells were isolated by FACS from 3 C57BL/6 mice (10 weeks-old 2 female, 1 male) and plated on the top of matrigel (7000-15000 cells per well). On day 9 the cells were starved for 22 hours and then treated for 6 hours with 50 ng/µl IL-1β (Peprotech, Cat# 211-11B), 2 µM Nutlin-3a (Selleck, Cat# S8059), 20 µM LPA or vehicle control as presented on the corresponding **Figure 1B**. On day 10, RNA was isolated from organoids and used for bulk RNA-seq. All organoids were dissociated from matrigel using cold PBS and 10 min incubation with 15U/ml dispase at 37°C, pelleted at 550 x g for 5 minutes at 4°C and harvested for RNA using method described in the section “RNA isolation”. RNA concentration was measured by Nanodrop. Freshly isolated RNA was stored at −80°C and sent on dry ice to Plasmidsaurus for sequencing and analysis. Library generation and sequencing were performed by Plasmidsaurus (Plasmidsaurus Inc.) using “3’ end counting method”. Single-end sequencing of ∼ 90 bp read length was performed on an Illumina platform at a read depth of at least ∼12 million reads per sample on average. Raw data (raw reads) in fastq.gz format was processed through a general pipeline, as described previously.^54^ Reads were aligned to the mm39 (GRCm39) mouse genome using Kallisto and imported into R Studio for analysis via the TxImport package. Transcripts were excluded from downstream analysis if they had zero counts across any sample or failed to reach a minimum expression threshold of one count per million in at least one sample. Differential gene expression analysis of raw read count data was performed using the DESeq2 package. We separately compared effect of each drug to control (n=3 for each sample). DEGs were exported and their overlapping was checked by using Venny 2.1^55^ as well as by Metascape^56^.

#### 2. Sample preparation for organoid bulk RNA-sequencing for short and long-term LPA treatment

FACS-sorted AT2 cells (Draq7-, CD45-, EpCAM+, Integrin β4-, Lysotracker green+) from 2 male and 2 female 8 weeks old wild-type C57Bl6 mice were seeded on top of 80µl Matrigel in organoids media with adding “passaging compound” only at initial plating. Later, media has been changed every 3 days, where regular organoid media was used for short-term groups and media with 10µM LPA or equivalent volume of ethanol was applied for long-term groups. Organoids were maintained in these conditions for 10 days. Detailed design of experiment is presented on the corresponding **Figure 2A**. For short-term treatment groups, at day 9 organoids were washed twice with 150µl DPBS and starved for 22 hours in 200µl of Advanced DMEM/F12. Then, organoids were incubated for 6 hours in media with 20µM LPA or equivalent volume of Ethanol. All organoids were dissociated from matrigel using cold PBS and 10 min incubation with 15U/ml dispase at 37°C, pelleted at 550 x g for 5 minutes at 4°C and harvested for RNA using method described in “RNA isolation”.

#### 3. Sample preparation for murine AT2s bulk RNA-sequencing

For this experiment aFos^+/+^; Sftpt^CreERT2+/+^ (10F, 10M, 6-9 weeks old) mice and were equally divided between 4 groups: PBS_Oil, PBS_Tamoxifen, Influenza_Oil, Influenza_Tamoxifen. To induce expression of aFos, aFos^+/+^; Sftpt^CreERT2+/+^ mice were treated with 0.25 mg/g of body weight tamoxifen by i.p. or oil as a control at days −25, −23, −21. After 3 weeks, mice were infected with influenza virus. At 14 days post-flu (dpi), AT2 cells were isolated and sorted by FACS (draq7-; CD45-; EpCAM+; Integrin β4-) for further analysis from 4 mice per group (2F, 2M). RNA from AT2 cells was isolated using the ReliaPrepTM RNA Cell Miniprep kit.

### Bulk RNA-sequencing by GENEWIZ

Quality and quantity of mRNA from chapters **“2”** and **“3”** were analyzed using a TapeStation (High Sensitivity RNA Screen Tape) and TapeStation Analysis Software 5.1 (Agilent Technologies). 1 ng of RNA was used to synthesize cDNA using Takara SMART-Seq HT Kit according to manufacture protocol and amplified for 15 cycles. Amplified cDNA was pulled by mixing with magnetic beads (1:1, AMPure XP, Beckman Coutlrt, Cat# A63880) and eluted in 17 µl of elution buffer (Takara, Cat# ST2098). Library generation and sequencing were performed by GENEWIZ Co. Ltd. Briefly, PolyA-selected RNA was used to generate libraries using the NEBNext Ultra II RNA Library Prep Kit for Illumina (NEB) according to the manufacturer’s instructions. Sequencing was performed using a 2 × 150bp paired-end configuration on an Illumnia HiSeq at a read depth of ∼45 million reads per sample on average. Raw data (raw reads) in fastq.gz format were downloaded and processed the same way as described in a chapter above. Reads were aligned to the mm39 (GRCm39) mouse genome using Kallisto and imported into R Studio for analysis via the TxImport package. Transcripts were excluded from downstream analysis if they had zero counts across any sample or failed to reach a minimum expression threshold of one count per million in at least one sample. Differential gene expression analysis of raw read count data was performed using the DESeq2 package. For **Figure 2**, we separately determined differentially expressed genes between short-term control and short-term LPA, as well as between long-term control and long-term LPA (n=4 for each sample). For **Figure 7**, we identified differentially expressed genes between uninfected, infected, aFos+, and WT groups. Each mouse represents one biological replicate. A single sample, “2016_F_aFos_Flu”, was excluded from analysis due to low *aFos* expression determined by RT-qPCR and low reads from sequencing.

### Analysis of publicly available data

VISIUM Spatial Transcriptomics dataset was previously generated in our lab and is deposited into the GEO under the accession code GEO: GSE326788 ^22^. For this study, no further analysis or modification have been done. Images and figures were generated using 10x Genomics Loupe Browser v7.0.1. Publicly available data of single cell RNA sequencing dataset generated by Gentile et al.^26^ and Niethamer et al. ^57^ were reanalyzed to generate dot plot using Seurat version 5.4.0 ^58^.

### Statistical Analysis

Data was collected in Microsoft Excel. Statistical analyses were performed using R and GraphPad Prism 7. Unpaired one-or two-tailed Student’s *t*-tests were used to determine statistical significance between two groups. For experiments with more than 2 groups, analysis of variance was performed. Details of sample size used statistical tests and definition of significance can be found at corresponding figure legends.

## Resource availability

### Lead contact

Requests for further information and resources should be directed to and will be fulfilled by the lead contact, Andrew E. Vaughan.

### Materials availability

All unique/stable reagents generated in this study are available from the lead contact, Andrew E. Vaughan, with a completed materials transfer agreement upon initial publication of this study.

### Data and code availability

All sequencing data and associated metadata are deposited in GEO (currently in progress). At the time of publication, all data generated in this study will be available from online repositories or can be obtained upon request from the corresponding author. This paper does not report original code.

## Acknowledgments

The authors appreciate all present and former Vaughan’s lab members (Dr. Joanna Wong and Dr. Gan Zhao) for their input in the project establishment and crucial critics. We thank the PennVet Transgenic Core and Center for Host-Microbial Interaction (Dr. Daniel Beiting and Daniel Cutillo) for their assistance in performing these studies. We also thank Dr. Jarod Zepp, Dr. Kate Hamilton and Dr. Nuala Meyer for invaluable insight into experimental design and serving on the T32 committee.

## Author Contributions

A.K.: designed the study, performed experiments, conducted analyses, wrote and edited the manuscript. A.I.W.: designed a mouse model. N.P.H., K.H.: performed experiments with human, conducted analyses. JK: provided human samples and supervised some experiments. S.K-G., M.M.M., M.M., E.A.M., D.M.A., M.Y.S.: helped to design the study, contributed to data analysis and/or editing this manuscript. A.E.V.: conceptualized and supervised this study, performed experiments, conducted analyses, wrote and edited the manuscript.

## Declaration of generative AI and AI-assisted technologies in the writing process

During the preparation of this manuscript, the authors used generative AI features built into Adobe Illustrator to create illustrations of cells, mice, lungs, and graphical abstract. No AI-tools were used to correct grammatical errors or improve language.

---

**Figure S1.**
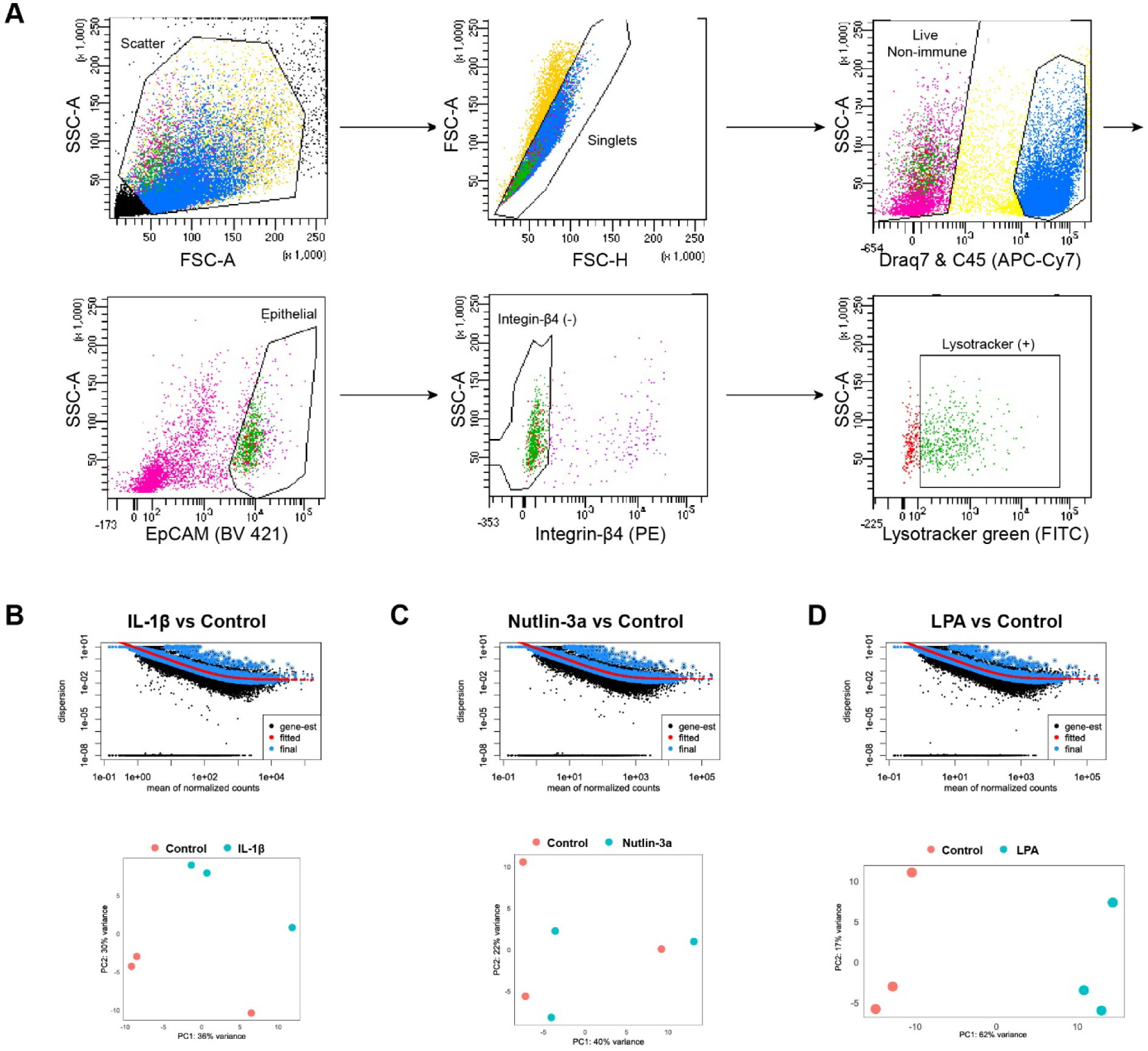
Gating strategy for isolation of murine AT2 cells by FACS. **(A)** Schematic showing representation AT2 FACS gating strategy up to Lysotracker green (-). All events were gated to single cells, viable cells (draq7 -), non-immune cell (CD45 -), epithelial cells (EpCAM +), Integrin-β4 (-). To determine Lysotracker green (+) gate, we included sample stained for all antibody but lacked lysotracker for each sorted biological replicate sample. **(B-D)** The dispersion estimates as a function of the mean of the normalized count (*top*) and PCA (*bottom*) for IL-1 β **(B)**, Nutlin-3a **(C)** and LPA **(D)** groups compared to control.

**Figure S2.**
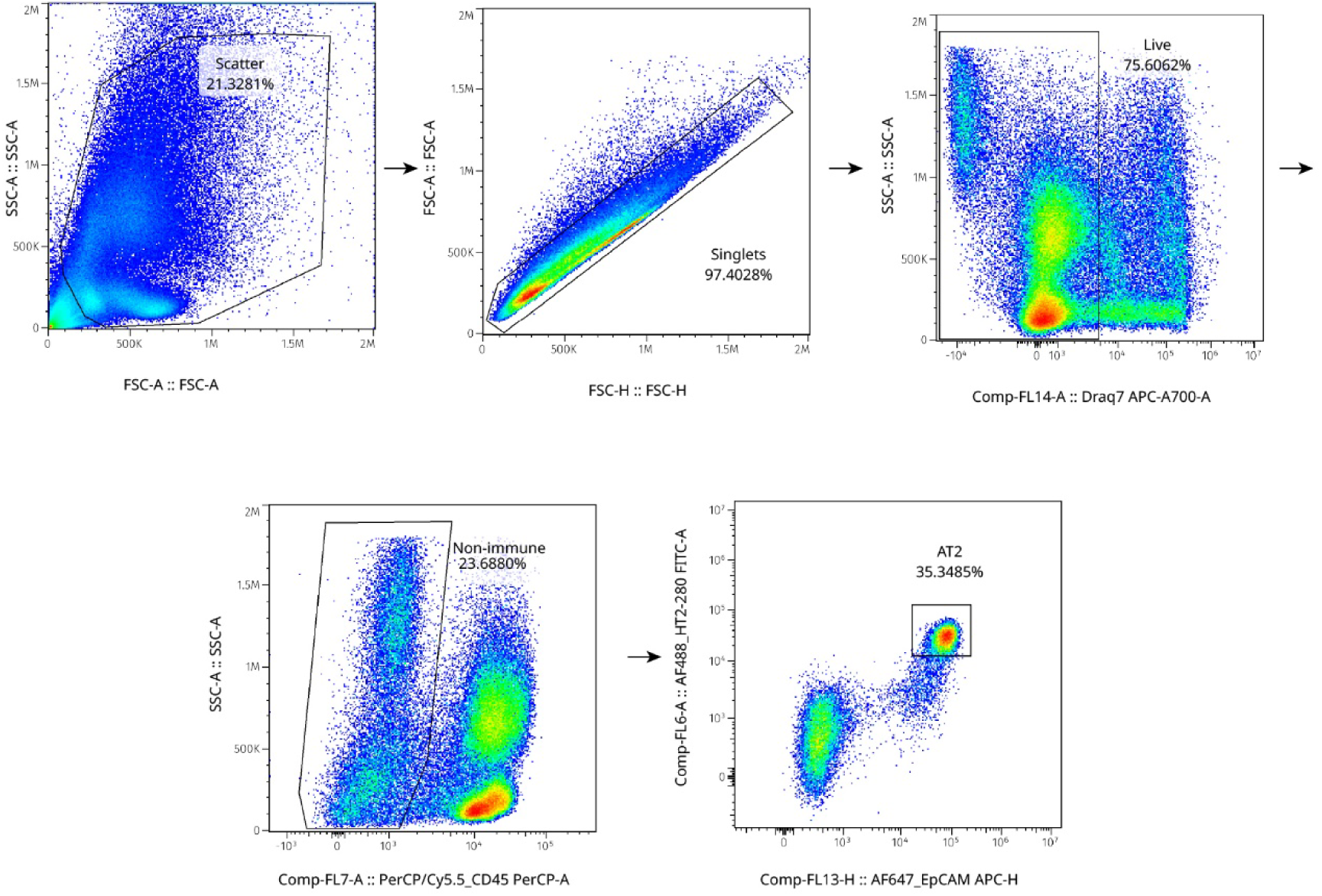
Gating strategy for isolation of human AT2 cells by FACS. Schematic showing representative human AT2 FACS gating strategy up to HT2-280. All events were gated to single cells, viable cells (draq7 -), non-immune cell (CD45 -), epithelial cells (EpCAM +), AT2s (HT2-280+).

**Figure S3.**
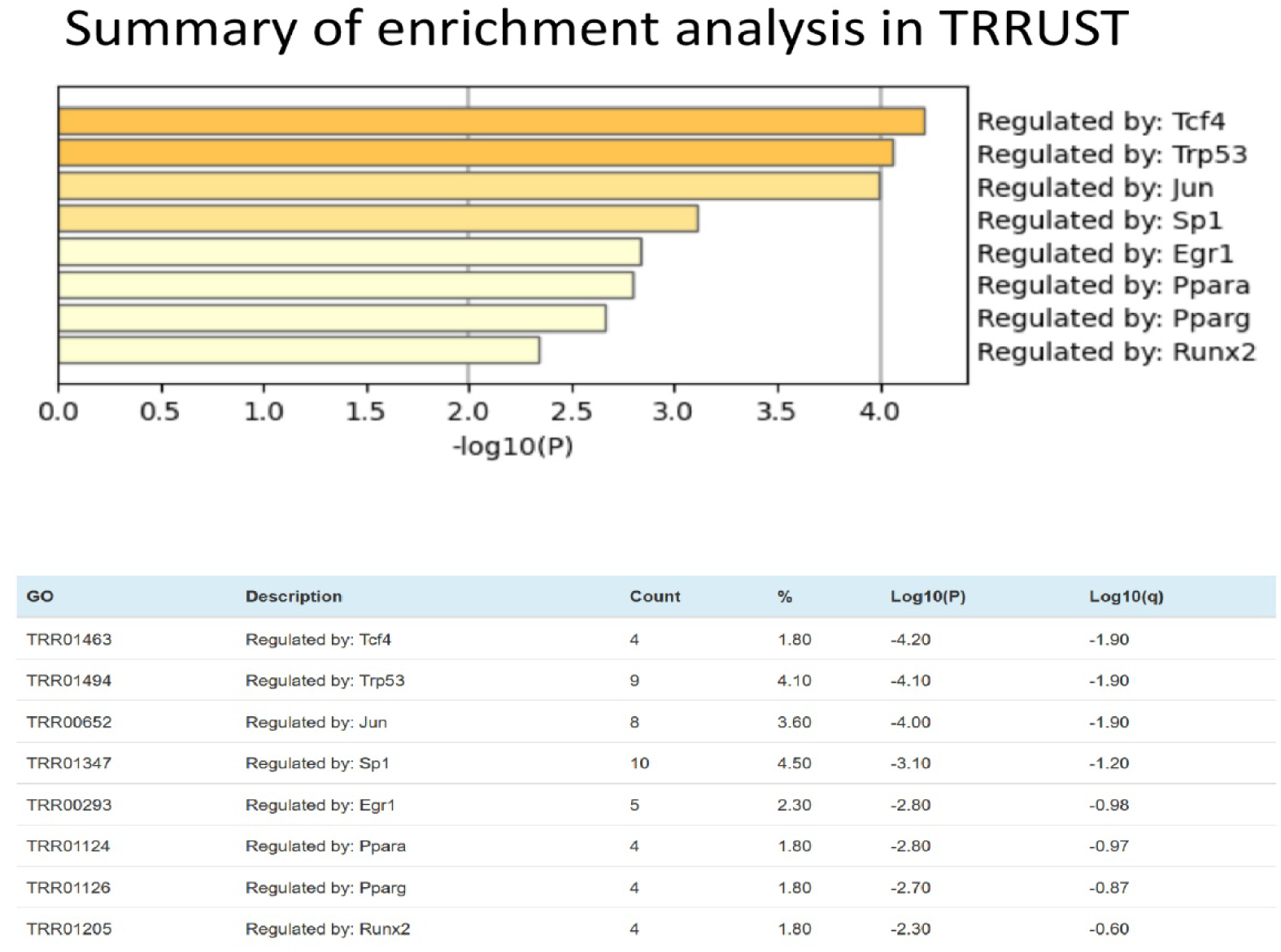
Analysis of activated transcription factors by Metascape on long-term LPA-treated group. Enrichment analysis of transcription factors (TRRUST database via Metascape) using the significantly upregulated genes in Figure 2H,**I** (long-term LPA vs long-term control).

**Figure S4.**
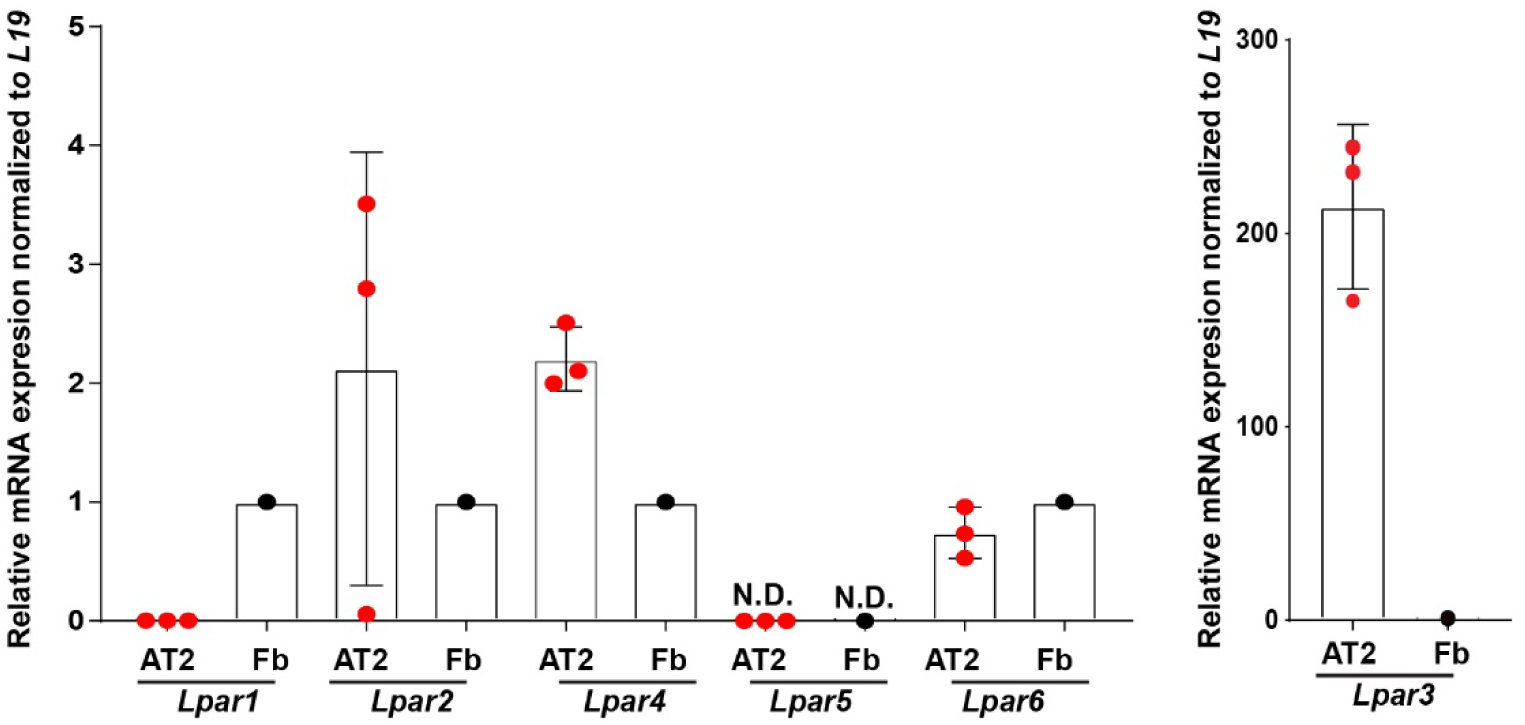
LPARs expression in murine AT2s and fibroblasts. *Lpar1-6* expression was validated by RT-qPCR in 100k freshly sorted primary murine AT2 cells and cultured murine fibroblasts (fb). Gene expressions are normalized to *L19* and to fibroblasts. Data shown as mean fold change ± SEM (n.d. – not determined). Each dot represents one biological replicate.

**Figure S5.**
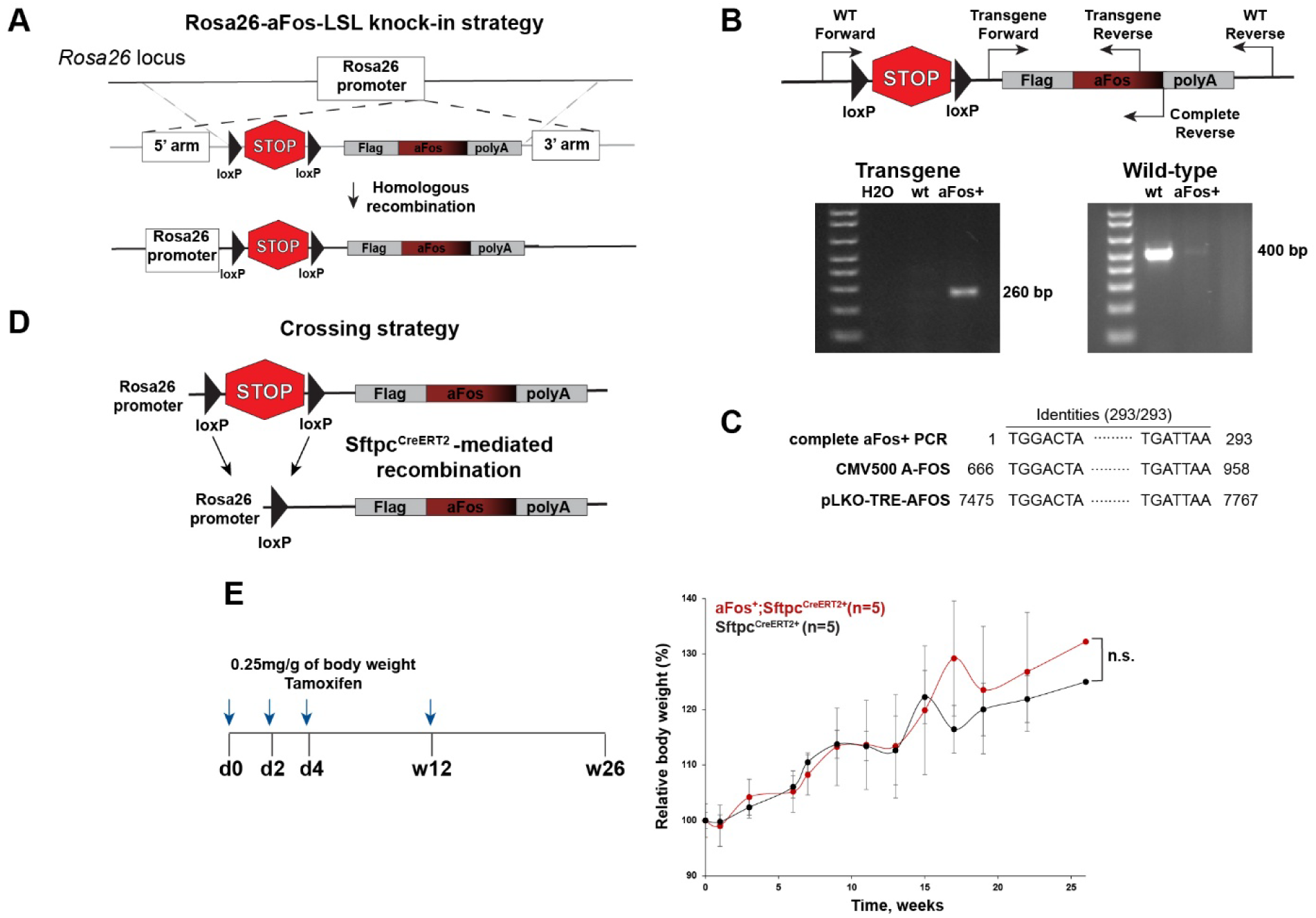
Generation and characterization of aFos-LSL mouse. **(A)** Schematic showing knock-in strategy for aFos-LSL mouse generation. To genetically inhibit activation of AP-1 signaling, we developed a mouse carrying the aFos construct on the Rosa26 locus with a stop codon placed prior to the transgene. These animals were then bred with mice carrying an AT2-specific CreERT2 (Sftpc^CreERT2+/+^). **(B)** Primer location and genotyping for transgene (*left*) and for wild-type allele (*right*). **(C)** Validation of aFos sequence. To ensure that established mouse line did not mutate over time, we isolated DNA from a murine tail after greater than 4 generations of breeding, amplified the full-length aFos construct and compared its sequence to original plasmid by using BLAST tool.^50^ Identity is 100% (293/293). **(D)** Mouse crossing strategy with the AT2-specific Sftpc-CreERT2 line. **(E)** Long-term effect of aFos expression. Experimental design (*left*). aFos^+/+^; Sftpt^CreERT2+^ and Sftpt^CreERT2+^ mice were treated with 0.25 mg/g of body weight tamoxifen by i.p. at days 0, 2, 4, and at week 12. Body weight curves (*right*). Graph showing changes in relative body weight day normalized to day 0 (n=5). Notably, 3 mice died immediately after i.p. injection performed at different time point, and they were excluded from further analysis. *P* values were calculated using an unpaired two-tailed t-test. No differences (n.s) were found between groups for both measurements: area under curve and a final body weight at the end point of experiment (week 26).

**Figure S6.**
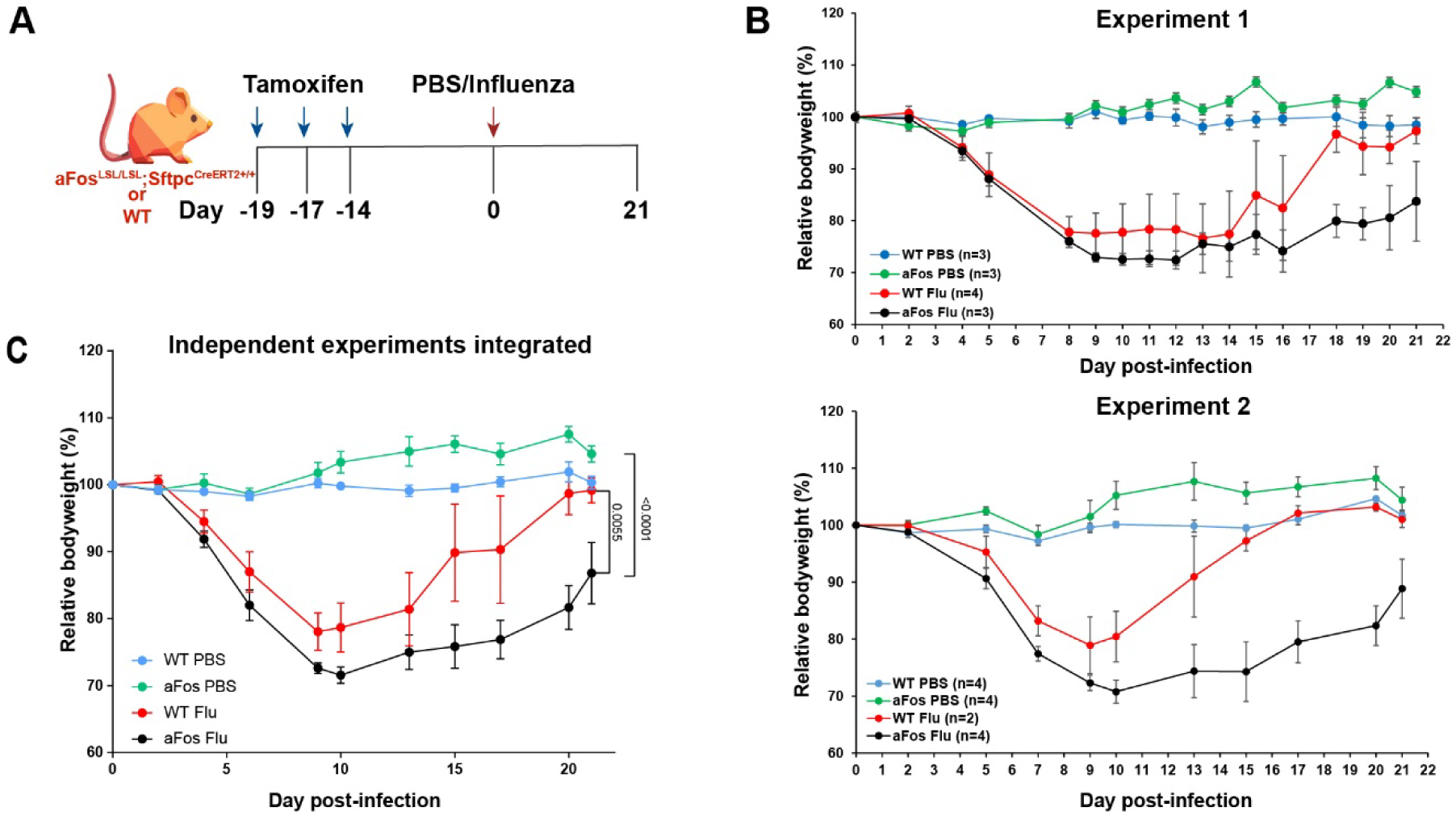
Impaired AP-1 signaling pathway in AT2s prevents normal recovery after influenza infection at least until day 21. **(A)** Schematic outline of the experimental design. aFos^+^; Sftpc^CreERT2+/+^ or functionally wild-type (wt, negative for at least one of the two transgenes) mice (4-8 weeks old, males and females) were treated with 0.25 mg/g of body weight tamoxifen by i.p. at days-19, −17, −14. After 14 days first injection, mice were infected with Influenza. Body weight was measured daily. We performed two independent experiments with similar experimental design. Only mice that lost at least 10% of initial body weight by day 6-9 were included in further analysis. **(B)** Body weight curves after infection for experiment 1 (*top*) and experiment 2 (*bottom*). Graph showing changes in relative body weight normalized to day 0 (mean ± SD).**(C)** Body weight curve for combined independent experiments. All mice from both experiments were included (n=6-7 per group). P values were calculated for day 21 using a two-way ANOVA with post-hoc Turkey multiple comparison test.

**Figure S7.**
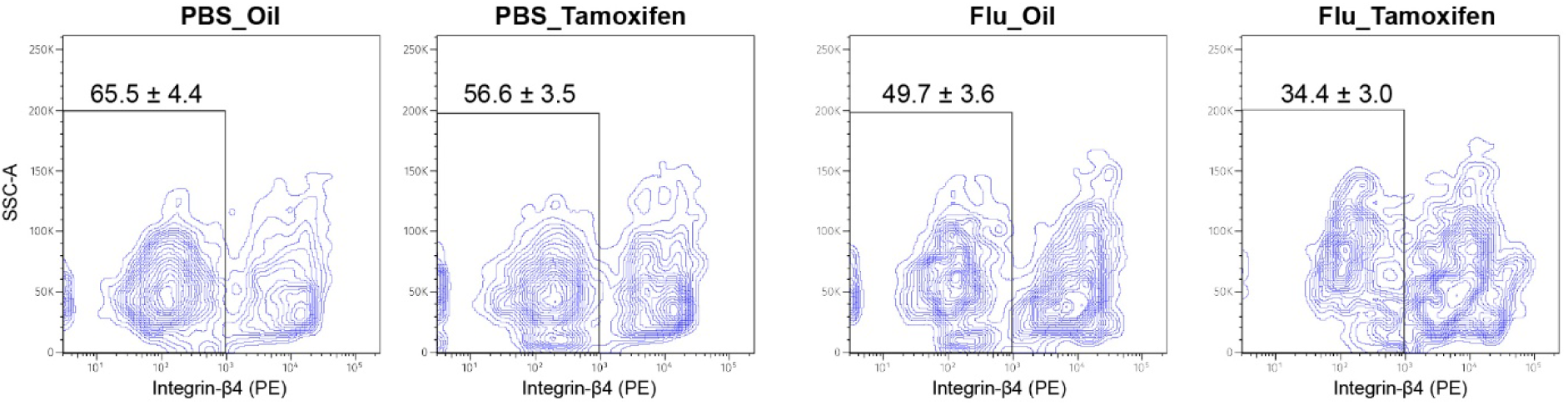
Representative gating scheme for identification of percentage of AT2 cells in murine lungs from the experiment presented on. Figure 6A. All events were gated to single cells, viable cells (draq7 -), non-immune cell (CD45 -), epithelial cells (EpCAM +) in the same way as it presented on **Figure S1**. AT2 cells were determined by the lack of Integrin-β4 expression. For this experiment only, Lysotracker was not used in the sort scheme because AT2s from injured lungs lose the lamellar bodies detected by Lysotracker. The same gates were applied across all samples. Data shows mean ± SD (n=4).

**Table S1.**
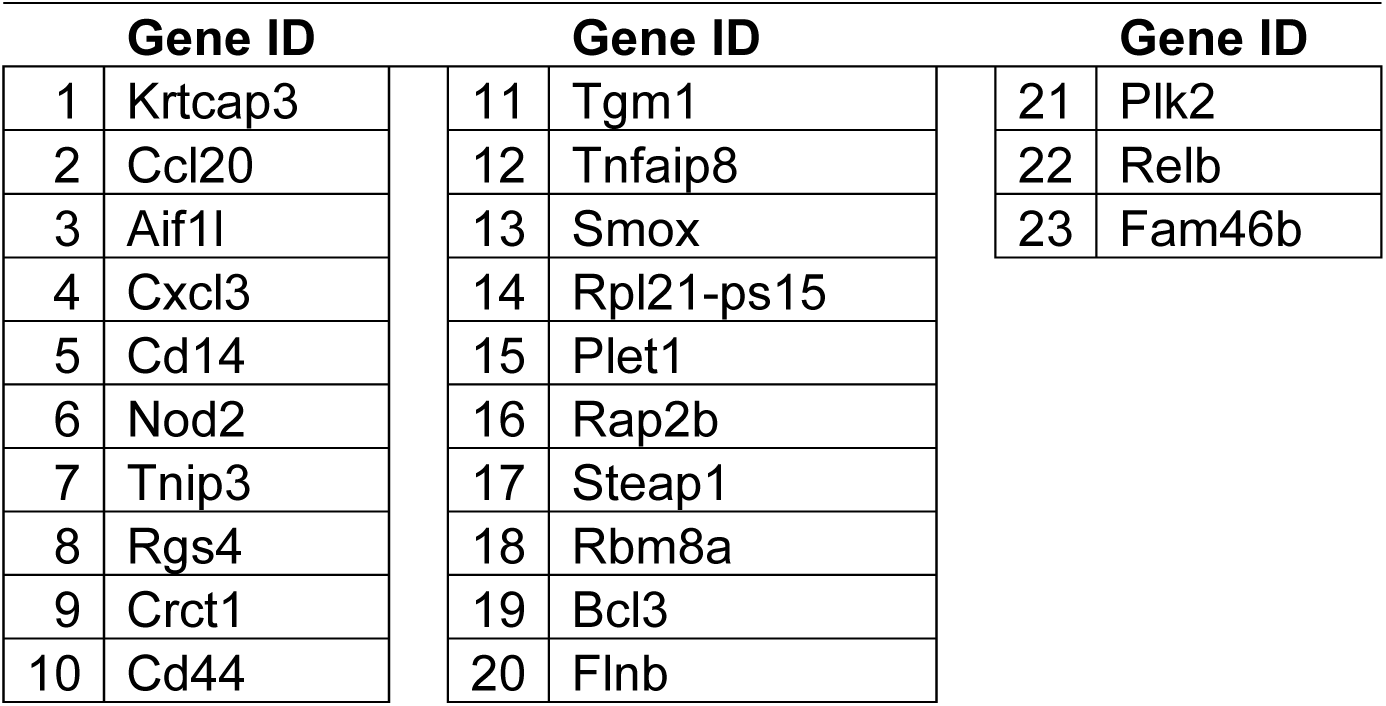
Gene list for 23 upregulated and overlap DEGs between IL-1β-and LPA-treated groups Gene ID Gene ID Gene ID. This dataset was used for prediction analysis for transcriptional factors for **Figure 1G**.

**Table S2.**
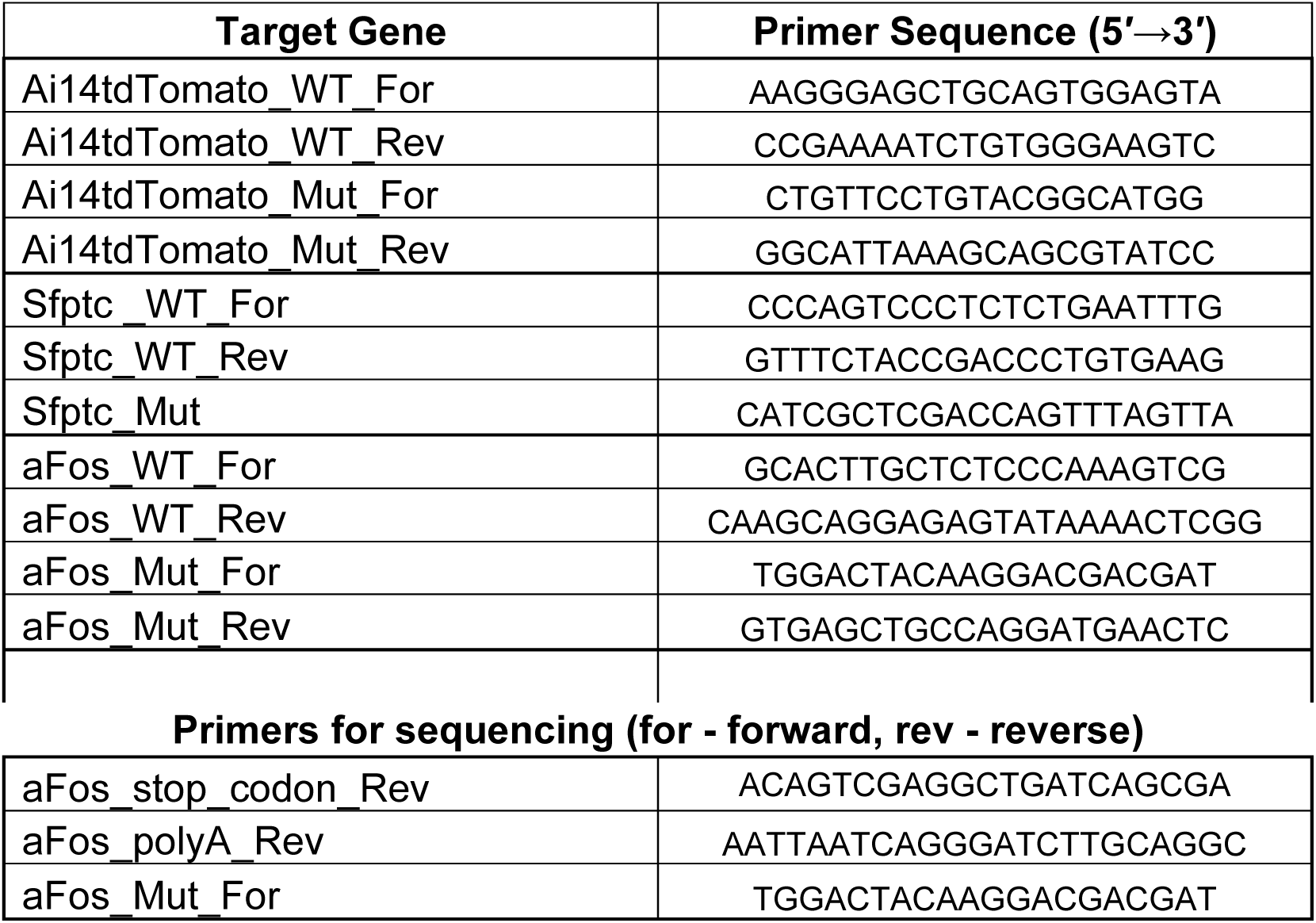
Primers for genotyping (for - forward, rev - reverse)

**Table S3.**
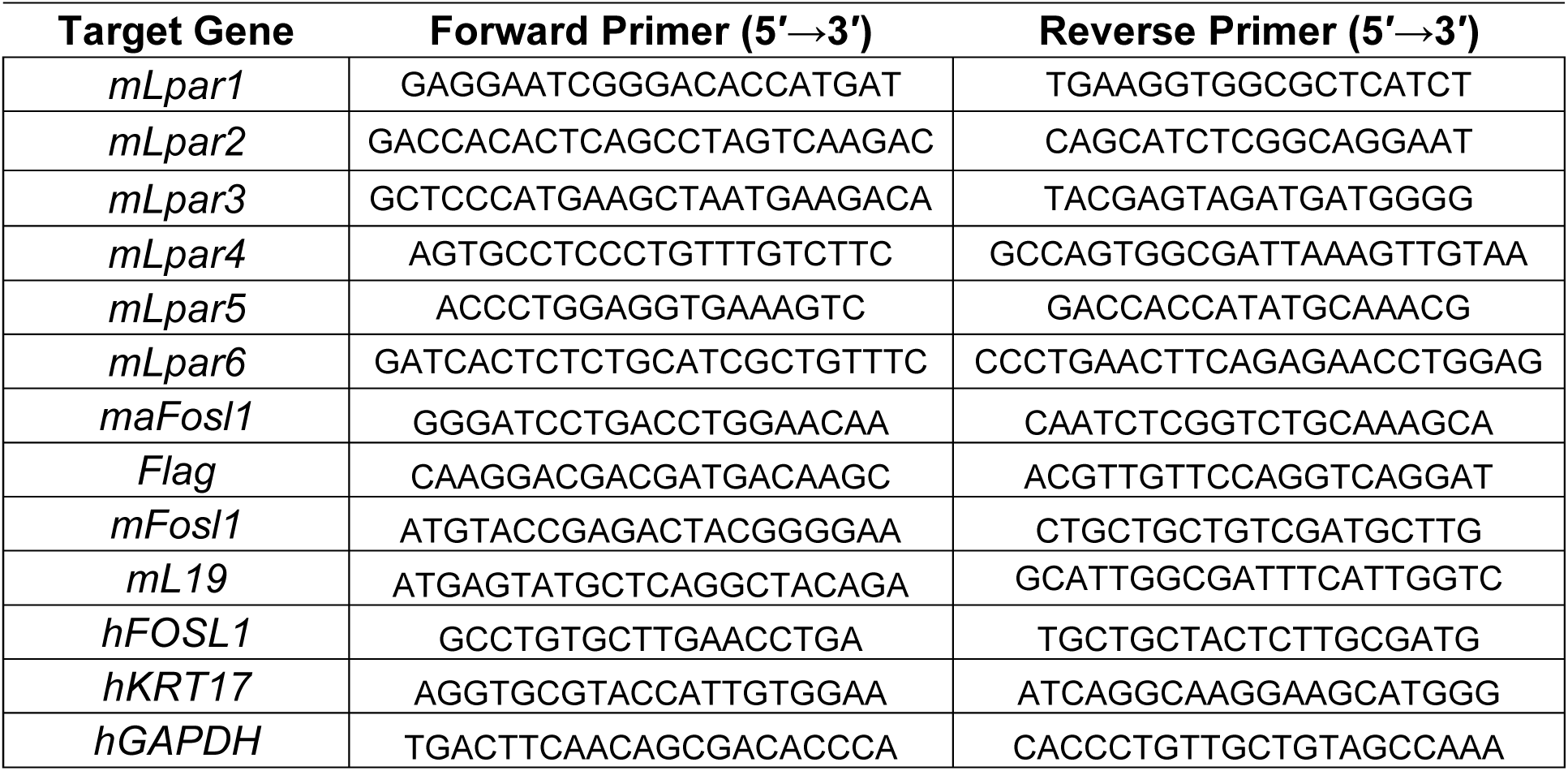
Primers for RT-qPCR (m - mouse, h - human)

